# From egg to adult: a developmental table of the ant *Monomorium pharaonis*

**DOI:** 10.1101/2020.12.22.423970

**Authors:** Arjuna Rajakumar, Luigi Pontieri, Ruyan Li, Rasmus Stenbak Larsen, Angelly Vásquez-Correa, Johanne KL Frandsen, Ab Matteen Rafiqi, Guojie Zhang, Ehab Abouheif

## Abstract

Ants are one of the most ecologically and evolutionarily successful groups of animals and exhibit a remarkable degree of phenotypic diversity. This success is largely attributed to the fact that all ants are eusocial and live in colonies with a reproductive division of labor between morphologically distinct queen and worker castes. Yet, despite over a century of studies on caste determination and evolution in ants, we lack a complete ontogenetic series from egg to adult for any ant species. We therefore present a developmental table for the Pharaoh ant *Monomorium pharaonis*, a species whose colonies simultaneously produce both reproductive queens and completely sterile workers. In total, *M*. *pharaonis* embryonic, larval, and pupal development lasts 45 days. During embryogenesis, the majority of developmental events are conserved between *M*. *pharaonis* and the fruit fly *Drosophila melanogaster*. However, we discovered two types of same-stage embryos prior to gastrulation: (1) embryos with internalized germ cells; and (2) embryos with germ cells outside of the blastoderm at the posterior pole. Moreover, we found two-types of embryos following germ band extension: (1) fertile embryos with primordial germ cells; and (2) sterile embryos with no germ cells. Together, our data shows that the queen (fertile) and worker (sterile) phenotypes are already determined and differentiated by early embryogenesis. During larval development, previous studies and our data find 3 larval instars reproductives and workers. However, there is considerable variation within each caste-specific instar, making it difficult to lineate instar boundaries. Here, we propose that developmental and anatomical markers can segregate larvae into gyne (unmatted queen), male and worker castes, including during the 1^st^ larval instar. Overall, we hope that the ontogenetic series we present here will serve as a blueprint for the generation of future ant developmental tables.

## Introduction

A major goal of evolutionary developmental biology (evo-devo) is to understand how developmental systems evolve and influence the evolution of organismal genotypes and phenotypes (Arthur 2002, Hall 2006, Carroll 2008, Hall 2012, Wagner 2014, Love 2014, Moczek et al. 2015). Over the last four decades, evo-devo has shown that the diversity in animal body plans involves tinkering with a relatively small number of highly conserved developmental regulatory genes, known as the ‘genetic toolkit’ (Patel et al. 1994, Quiring et al. 1994, Caroll 1995, Gerhart and Kirschner 1997, Akam 1998, Hall 2003, Carroll 2005, Davidson and Erwin 2006, Peter and Davidson 2016, Hu et al. 2019, Bruce and Patel 2020, Murugesan et al. 2022). Evo-devo studies on ants have played a crucial role in revealing the ecological dimensions of *evo*-*devo*, a re-emerging synthesis known as ‘*eco*-*evo*-*devo*’ (Metzl et al. 2018). This field explores how the interaction between this highly conserved genetic toolkits and environmental factors, such as nutrition, temperature, and social interactions can generate or mask phenotypic variation that may facilitate or constrain evolutionary change (Evans and Wheeler 1999, Miura et al. 1999, West-Eberhard 2003, Abouheif et al. 2014, Gilbert et al. 2015, Santos et al. 2015, Sommer and Mayer 2015, Toth and Rehan 2017, Rajakumar and Sanger 2018, Kapheim et al. 2020).

Ants are particularly exciting models for *eco*-*evo*-*devo* because all ants are eusocial. Eusociality is the highest-level of social organization where individuals have obligate colonial lifecycles. Notably, individuals within a colony bear dramatically different phenotypes called “castes”, typically queen and workers, to divide labour. Castes within a single colony are polyphenic, meaning that the same genome can initiate a queen or a worker developmental program in response to environmental cues (Brian 1974, Passera and Suzzoni 1979, Hölldobler and Wilson 1990, Hölldobler and Wilson 2009, Penick et al. 2012). In most ant species, queen and worker castes display dramatic differences in morphology and life history. Key differences include wing polyphenism, where colonies develop winged queens and wingless workers; a size and reproductive asymmetry between hyperfertile queen and subfertile worker castes; and within the worker caste, the evolution of a novel soldier subcaste. Each of these innovations evolved through the differential regulation of the genetic toolkit during either embryonic, larval, or pupal stages (Abouheif and Wray 2002, Wheeler and Nijhout 2003, Sameshima et al. 2004, Gotoh et al. 2005, Khila and Abouheif 2008, Khila and Abouheif 2010, Rajakumar et al. 2012, Gotoh et al. 2016, Béhague et al. 2018, Rajakumar et al. 2018, Oettler et al. 2019, Yan et al. 2022, Brahma 2023). Furthermore, within ant societies, developing larvae play fundamental roles in social regulation, such as in processing food, producing pheromones and regulating caste-ratio, further increase the complex regulatory landscape of colonial life (Mamsch 1967, Villalta et al. 2015, Ebie et al. 2015, Warner et al. 2016, Schultner et al. 2017, Warner et al. 2019, Snir et al. 2022).

To date, empirical eco-evo-devo studies on ants have helped us to understand queen-worker caste differentiation (Wheeler 1986, Abouheif and Wray 2002, Sameshima et al. 2004, Gotoh et al. 2005, Khila and Abouheif 2008, Khila and Abouheif 2010, Miyazaki et al. 2010, Penick et al. 2012, Qui et al. 2018, Chandra et al. 2018, Warner et al. 2019, Nagel et al. 2020, Qui et al. 2022, Yan et al. 2022, Trible et al. 2023), epigenetic regulation (Alvarado et al. 2015, LeBoeuf et al. 2016, Simola et al. 2016, Glastad et al. 2021, Gospocic et al. 2021), worker polymorphism (Wheeler and Nijhout 1981a, Wheeler and Nijhout 1981b, Rajakumar et al. 2018, Klein et al. 2018), social behaviour (Teseo et al. 2014, Trible et al. 2017, Yan et al. 2017, Glastad et al. 2021, Gospocic et al. 2021, Brahma et al. 2023, Ju et al. 2023), modularity (Yang and Abouheif 2011, Londe et al. 2015, Hart et al. 2024), evolutionary novelty (Rajakumar et al. 2012, Favé et al. 2015, Rajakumar et al. 2018, Powell et al. 2020, Rafiqi et al. 2020), gene by environment interactions (Schrader et al 2014, Singh and Linksvayer 2020, Schrader et al. 2021, Glastad et al. 2023), ancestral developmental potentials (Rajakumar et al. 2012), and major evolutionary transitions in individuality (Bernadou et al. 2018, Rafiqi et al. 2020). Despite the success of these eco-evo-devo studies in ants, there is a general absence of a detailed ontogenetic series from egg to adult for any ant species. This has hampered our ability to advance the functional characterization of the genes and gene networks underlying key innovations in ants, such those regulating caste determination between queens and workers. For example, the ability to establish transgenic lines with CRISPR-CAS9 gene editing in any organism is predicated on a working knowledge of the timing and stages of that organism’s development. This is evident in recent studies that have functionally manipulated gene expression during development in different ant lineages, where an understanding of developmental timing was required to ensure the specificity and efficacy of reagents to downregulate gene expression (Alvarado et al. 2015, Trible et al. 2017, Yan et al. 2017, Rajakumar et al. 2018, Rafiqi et al 2020, Gospocic et al. 2021, Qui et al. 2022 Yan et al. 2022, Glastad 2023, Ju et al. 2023, Hart et al. 2023).

The general lack of developmental tables in ants is surprising given the long and rich history of the study of ant development. Prior to the rise of inclusive fitness theory (Hamilton 1964a and b), a group of biologists generated a body of pioneering work on the eco-evo-devo of ants (Metzl et al. 2018). This pioneering work includes studies by W.M. Wheeler (Wheeler 1893, Wheeler 1910, Wheeler 1911), Goestch (Goestch 1937, 1939), Bier (Bier 1952), and Dewitz (Dewitz 1878), all of whom made prescient insights into the developmental basis of caste determination. Furthermore, work by Ganin, (Ganin 18<u>6</u>9) Tanquary (Tanquary 1912), Blochmann (Blochmann 1892), Strindberg (Strindberg 1913, 1914, 1915, 1916, 1917) Hegner (Hegner 1915), Lilienstern (Lilienstern 1932), and Buchner (Buchner 1918, Buchner 1965) began investigating, in remarkable detail, ant embryogenesis (Figure 1). To our knowledge, Ganin is one of the first to present an ontogenetic series of ant embryonic development, in which he described the embryonic development of *Formica fusca*, from the syncytial blastoderm stage through to segmentation (Ganin 1869). Later, W.M. Wheeler in 1918 and 1922 (Wheeler 1918, Wheeler 1922) described the general morphology of ant larvae, and later in 1947, Athias-Henriot provided the first detailed account of the internal anatomy of larvae (Athias-Henriot 1947). Finally, George Wheeler and Jeanette Wheeler, who in a long series of publications between 1953 and 1990 extensively described the external morphology of larvae of approximately 800 species of ants (Wheeler and Wheeler 1953, Wheeler and Wheeler 1955, Wheeler and Wheeler 1976, Wheeler and Wheeler 1990). Only more recently have researchers begun to describe in detail the number of instars in a given ant species (Passera 1971, Passera 1973, O’neal and Markin 1975, Petralia 1979, Wheeler 1983, Sameshima et al. 2004, Fox et al. 2012, Fox et al. 2017, Masuko 2017, Solis et al. 2010; Alvarado et al. 2015, Koch et al. 2021).

**Figure 1.**
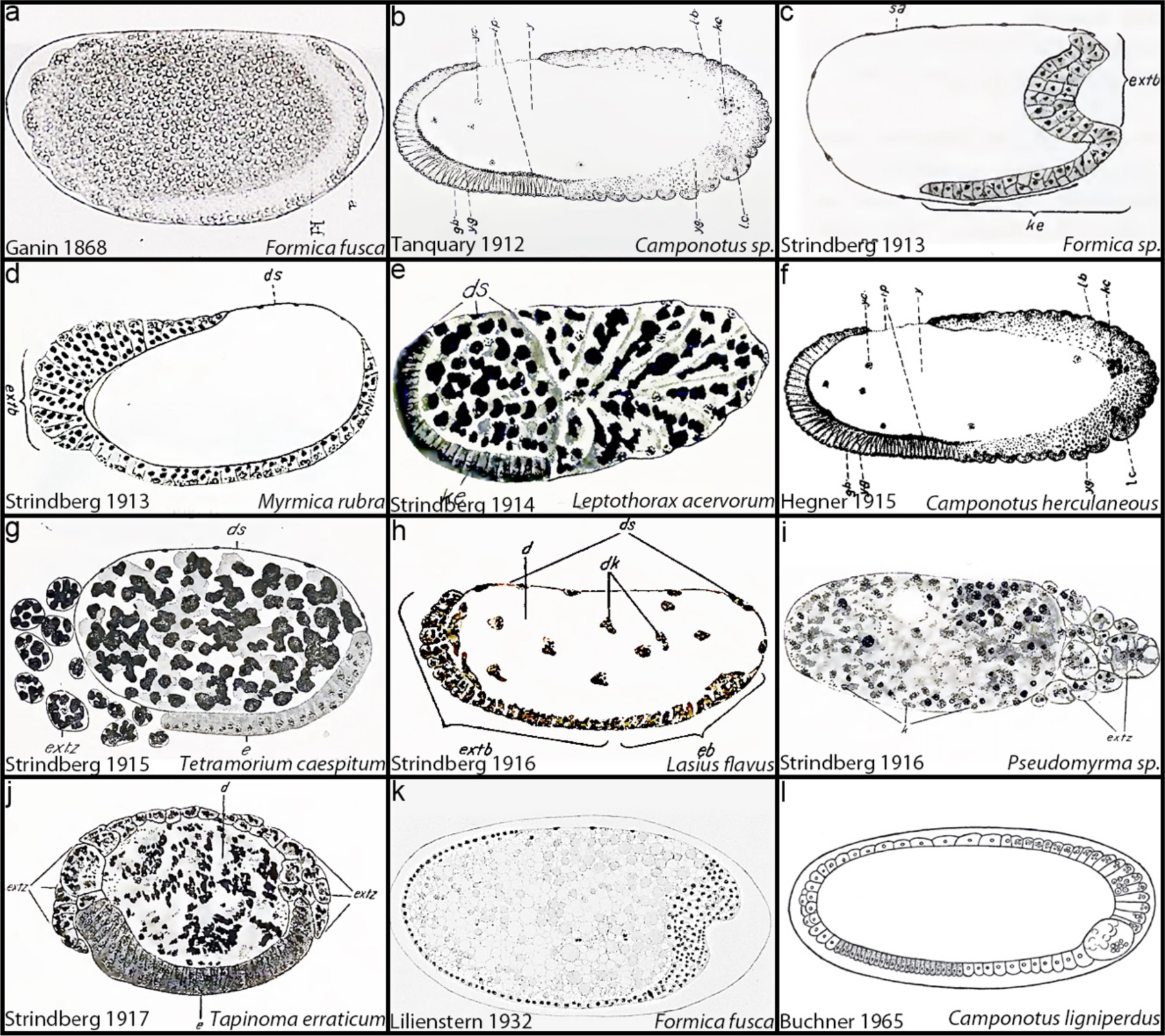
A century of ant embryology. a-l) Histological sections of cellular blastoderm or gastrulation stage embryos representing 4 ant subfamilies adapted from historical references. Embryos are in chronological order of their description. a) *Formica fusca* embryo adapted from Ganin 1868. b) *Camponotus sp*. embryo adapted from Tanquary 1912. c) *Formica sp*. embryo adapted from Strindberg 1913. d) *Myrmica rubra* embryo adapted from Stindberg 1913. e) *Leptothorax acervorum* embryo adapted from Strindberg 1914. f) *Camponotus herculaneous* embryo adapted from Hegner 1915. g) *Tetramorium caespitum* embryo adapted from Strindberg 1915. h) *Lasius flavus* embryo adapted from Strindberg 1916. i) *Pseudomyrma sp*. embryo from Strindberg 1916. j) *Tapinoma erraticum* embryo adapted from Strindberg 1917. k) *F*. *fusca* embryo adapted from Lilienstern 1932. l) *Camponotus ligniperdus* adapted from Buchner 1965. Anterior is to the left, posterior to the right, dorsal up, ventral down, except e and h where posterior is to the left and anterior to the right. Panels are ordered historically from oldest to latest.

Here we characterize an ontogenetic series, spanning embryonic, larval, and pupal stages of the Pharaoh ant *M. pharaonis*, a global invasive species. *M. pharaonis* colonies are polygynous (multiple queens), monoandrous (singly-mated), and have a monomorphic worker caste (limited size variation) (Jackson et al. 2004). Over the last decade, *M*. *pharaonis* has been used as a model to study the sociogenomic basis of social insect castes (Pontieri et al. 2017, Qui et al. 2018 Walsh et al. 2018, Warner et al. 2019, Walsh et al. 2020, Singh and Linksvayer 2020, Nagel et al. 2020, Li et al. 2022, Qui et al. 2022), collective behaviors (Gordon 2019, Walsh et al. 2020), and caste/sex ratio regulation (Warner et al. 2016, Pontieri et al. 2017,Warner et al. 2018, Singh and Linksvayer 2020). *M*. *pharaonis* is also a promising model for studying gene-environment interactions during development and evolution because reproductive (gynes and males– where a gyne is an unmatted queen) practice within nest mating, and workers are obligately sterile, lacking a germline. As such, only queen can produce brood within a colony (Hölldobler and Wilson 1990). These features make it possible to establish and maintain colony lineages, or ‘sociogenetic lines’ for several generations in the lab (Walsh et al. 2020). Furthermore, reproductives can be mated with unrelated partners, allowing the establishment of genetically heterogeneous crossed lines (Schmidt et al. 2011, Pontieri et al. 2017). Finally, *M. pharaonis* is particularly suited for addressing the question of caste determination and differentiation. Unlike the majority of ant species where queens produce gynes and males during only a short period of the year, making it difficult to pinpoint the contrasting developmental trajectories of gyne-destined and worker-destined brood. *M*. *pharaonis* produces all castes and sexes year-round, thereby enabling the ability to study the developmental programs of all castes at the same time (i.e., worker, gyne, male (Edwards 1987, Qui et al. 2022). Altogether, a developmental table of *M. pharaonis* would greatly facilitate all these areas of study, especially the eco-evo-devo of ants.

## Materials and Methods

### Ant colony maintenance and collection of eggs, larvae and pre-pupae

Two colonies (D03 and 4030) of *M*. *pharaonis* were used in this study to collect egg, larval, pre-pupal and pupal stages. This colony was part of a larger stock of colonies artificially created in 2010 through the sequential cross of eight inbred lineages (see Schmidt et al. 2011 and Pontieri et al. 2017 for breeding methods) and has since been maintained at the University of Copenhagen. The colony was kept at 27 ± 1 °C and 50% RH in a plastic box (27 × 17 × 9.5 cm) coated with Fluon® (polytetrafluorethylene, De Monchy, The Netherlands), with cotton-sealed plastic tubes serving as nesting sites. The colony was fed twice a week with a standardized diet containing a 1:4 ratio of total proteins to digestible carbohydrates (diet modified from Dussutour and Simpson 2008) and house crickets (*Acheta domesticus*, QB Insects, Linnich, Germany). Water was provided *ad libitum*.

For timed egg collections, 15 queens were removed from the colony using a fine brush and placed in a Fluon®-coated petri dish (15 cm x 1.5 cm) containing a 2 cm^2^ piece of black cardboard to nest underneath. Queens were provided with 80 workers from their colony, a piece of food and a water tube. Queens were left to lay eggs for different intervals in order to determine how long a given embryonic stage lasts. Egg depositions were either collected at the same time as queen removal (for stages 1 and 2) or aged in the petri dish (for stages 3 and onwards), with workers tending them (see table 1). Food replaced daily, and petri dishes containing aging eggs were kept at the same temperature and humidity condition as the stock colony. Eggs were then collected at the desired stage and fixed.

**Table 1.** Sampling scheme of *M. pharaonis* embryos for staging. Left column: Time intervals that eggs were collected following the isolation of *M*. *pharaonis* queens. Middle column: The number of embryos fixed, DAPI stained and staged at a given time period. Right column: The stages represented during this time window.

| Time interval (hours) | Number of embryos | Stages identified |
| --- | --- | --- |
| 0-6 | 32 | 1 |
| 6-12 | 32 | 2 |
| 12-18 | 41 | 2, 3 |
| 18-24 | 28 | 3, 4 |
| 24-27 | 17 | 4, 5 |
| 24-30 | 137 | 4, 5 |
| 27-30 | 34 | 5 |
| 30-33 | 16 | 5 |
| 30-36 | 40 | 5 |
| 36-42 | 77 | 6 |
| 42-48 | 120 | 7 |
| 48-54 | 24 | 8 |
| 54-60 | 10 | 8 |
| 60-66 | 39 | 8 |
| 66-72 | 40 | 9 |
| 72-84 | 44 | 9, 10 |
| 84-96 | 50 | 10 |
| 96-108 | 83 | 11 |
| 108-120 | 27 | 11 |
| 120-132 | 19 | 11 |
| 132-144 | 78 | 11 |
| 144-156 | 55 | 11 |
| 156-168 | 48 | 11 |
| 168-180 | 46 | 12 |
| 180-192 | 31 | 12 |
| 192-204 | 16 | 13 |
| 204-216 | 14 | 14, 15 |
| 228-240 | 28 | 16, 17 |
| **Total** | **1226** |  |

Larval stages used for morphometric analyses were collected from two sub-colonies established by splitting in half the colony used for egg depositions. Previous studies have shown that the removal of queens allows workers to raise existing reproductive brood to adulthood (Edward 1987). Therefore, we created to sub-colonies to facilitate the collection of worker-destined and reproductive-destined brood. The first sub-colony, containing approximately 50 queens, was used to collect 1^st^ larval instars of unknown caste, 2^nd^ and 3^rd^ worker larval instars. The second sub-colony, dequeened at the time of its establishment in order to trigger the production of new reproductives (gynes and males), was used to collect 1^st^ larval instars of unknown caste, 2^nd^ and 3^rd^ reproductive larval instars. Individual larvae and pre-pupae were collected using a fine brush and gently lined up, with the cephalic capsule facing up, on a piece of double-side tape which was then placed on a microscope slide for imaging.

### Embryo fixation and nuclear staining

Embryos were gently transferred with a moistened brush in an incubation basket with a 100 µm mesh (Intavis Bioanalytical Instruments AG), dechorionated in 25% commercial bleach (from a 12.5% sodium hypochlorite stock) for 2 minutes and quickly washed under demineralized tap water for 30 seconds. Embryos were then bathed in 0.3% PBTx (1 x PBS, Triton X-100 – Sigma Aldrich) for 5 minutes on ice, heat fixed by boiling for 30 seconds in 0.3% PBTx, quickly quenched in 1 x PBS on ice and finally bathed in 0.1% PBTw (1 x PBS, Tween-20 – Sigma Aldrich) on ice for 5 minutes. Embryos were then washed 4 times in PBTw (5 minutes each wash), fixed in 4% paraformaldehyde for 30 minutes at room temperature, and then washed 3 times in 1 x PBS. Fixed embryos were transferred from the incubation basket to a 2 ml screw top clear glass vial (Supelco; Sigma-Aldrich) using a glass Pasteur pipette. After removing the 1 x PBS, 500 µl of ice-cold methanol were added and the vial was vigorously shaken for 10 seconds. Embryos were finally washed 2 times with ice cold methanol and either stored at −20 °C or immediately rehydrated. While, in other insects this methanol shock helps to crack the vitelline membrane or remove it entirely (Rafiqi et al. 2011), in *M. pharaonis* we found ice cold methanol did not remove the vitelline membrane but instead generated a larger separation between vitelline membrane and the embryo which increased image quality. Prior to their use in downstream staining, embryos were rehydrated through a series of methanol/PBTw dilutions (75%, 50%, 25%), and finally washed 3 times in PBTw. After removing the PBTw, embryos were incubated with a single drop of VectaShield mounting medium with DAPI (Vector Laboratories) for 10 minutes at 4 °C in dark. Fixed DAPI counterstained embryos were transferred along with the mounting media onto a microscope slide, covered with a cover slip and imaged.

### Hybridization Chain Reaction

For embryonic florescent *in situ* hybridization experiments, embryos were not treated with bleach during the fixation process. Instead, the chorion was manually removed and the vitelline membrane was left intact for pre-gastrulation stage embryos. For embryos after gastrulation, the vitelline membrane was removed manually. *M. pharaonis nanos and oskar* mRNA were visualized using the Hybridization Chain Reaction (HCR) method (Choi et al., 2018; Molecular Instruments). For *nanos*, 20 probes were generated against *M. pharaonis nanos* (XM_012677141). For *oskar*, 20 probes were generated against *M. pharaonis nanos* (XM_012677375). Hybridization Chain Reaction was performed using the manufacturers buffers and protocols (Choi et al., 2018; Molecular Instruments).

For wholemount fluorescent *in situ* hybridization of larvae HCR was performed following the whole-mount drosophila HCR v3.0 protocol (Choi et al., 2018) with some modifications. *M. pharaonis* larvae were fixed at room temperature in scintillation vials with 50% FPE (4% formaldehyde; 0.5× PBS; 25 mM EGTA) and 50% heptane. Fixation time was then adjusted so that 1^st^ and 2^nd^ larval instars were fixed for 2-3 hours, and 3^rd^ larval instars were fixed for 12 hours. Following fixation, the lower layer (FPE) was removed and replaced with methanol followed by vigorous shaking. The lower layer was replaced once more with methanol, at which point larvae sink to the bottom of the vial. Larvae were then dehydrated with several changes of methanol and stored at −20 °C. Proteinase K concentration and treatment time was adjusted to 50 μg/mL for 7 minutes for 1^st^ and 2^nd^ larval instars and 60 μg/mL for 7 minutes for 3^rd^ larval instars. Following amplification, one SSCT wash (5X SSC; 0.1% Tween-20; pH 7.0) was extended overnight with the addition of SYTOX™ Deep Red (1:1000) for nuclear staining in 1^st^, 2^nd^ and 3^rd^ larval instars. For *vasa*, 20 probes were generated against *M*. *pharaonis vasa* (XM_012686851.3). For *headcase*, 20 probes were generated against *M*. *pharaonis headcase* (XM_036286481.1).

### Antibody staining

Cellular blastoderm embryos were fixed and dissected as discussed above. Antibody staining was done as described in Rafiqi et al. 2020. Rabbit anti-Vasa antibody (gift from Paul Lasko) was used at a concentration of 1:100. Anti-rabbit Alex Flour 555 secondary antibody (Cell signaling, #4413) was used at a concertation of 1:500.

**Table 2.** Sample size of *M. pharaonis* embryos for a given stage. Left column: Stage classified after DAPI staining. Right column: Total number of embryos we examined of a given stage.

| Stage | Total number |
| --- | --- |
| 1 | 32 |
| 2 | 73 |
| 3 | 69 |
| 4 | 182 |
| 5 | 244 |
| 6 | 77 |
| 7 | 120 |
| 8 | 73 |
| 9 | 84 |
| 10 | 94 |
| 11 | 310 |
| 12 | 46 |
| 13 | 16 |
| 14 | 14 |
| 15 | 14 |
| 16 | 28 |
| 17 | 28 |

### Imaging

For embryos, DAPI images were captured at 20x magnification on an upright Olympus BX63F fluorescence microscope (Tokyo, Japan) equipped with a Retiga 6000 camera system (Qimaging, Surrey, British Columbia, Canada) using cellSens Dimension software version 1.16. For each embryo, a series of Z-stack images were acquired (step), going from one side to the other. Z-stack images were then deconvolved (Gaussian) and only relevant stacks were selected to create either a maximum or an average intensity projection z-stack image. Images were further processed in Adobe Photoshop CC to optimize brightness and contrast.

For brightfield images and videos of embryos, eggs were collected from the stock colony using a fine brush, placed on a microscope slide and gently covered with a cover slip. Drops of halocarbon oil 700 (Sigma-Aldrich) were released on the side of the cover slip and capillary action allowed the oil too slowly displace air and submerge the eggs. Eggs were not dechorionated. Lastly, for brightfield embryo images (Figure 4), embryos were processed in photoshop by changing brightness and contrast, along with sharpening the image, which allowed visualization of cell contours within the blastoderm. Eggs were imaged using the same protocol described for DAPI counterstained embryos. For videos, pictures were taken at different focal points every 20 minutes at room temperature.

For HCR-stained embryos, embryos were imaged using a Leica SP8 confocal microscope. Images are presented as maximum intensity Z-stacks, which were compiled using ImageJ2 (Rueden et al. 2017).

For brightfield images of whole larvae to make the images less distracting we subtracted the background to make the larvae over a black background (Figure 6).

For wholemount HCR-stained larvae, larvae were transferred into increasing concentrations of glycerol in 5X SSC and mounted in ProLong™ Glass Antifade Mountant for imaging. Images were captured on a Leica STELLARIS 8 inverted confocal laser scanning microscope. Image stacks were processed using Fiji/ImageJ (Rueden et al. 2017).

For imaging of pupae, pupal developmental series were obtained by isolating pre-pupae stages of reproductives and workers in Fluon® coated petri dishes. As soon as the individuals pupated, they were imaged on consecutive days using the same equipment and procedure as described for larvae and pre-pupae image acquisition.

### Larvae and pre-pupae measurements

Larvae and pre-pupae images were acquired using a BK plus lab system (Dun, Inc, Virginia, USA) equipped with a Canon 7D camera. Z-stack images were taken for each individual and combined using the software Zerene Stacker Professional edition. Adobe Photoshop CC was used to measure the maximum cephalic capsule and larval length. Larvae for initial instar identification were collected based on morphological traits previously used in the literature for instar classification (hair types and their presence/absence; mandible coloration).

In total, we collected 340 individuals, which were classified as follow: 103 1^st^ instars (62 from queenless, 41 from queenright), 51 2^nd^ instar workers, 55 3^rd^ instar workers, 35 pre-pupa workers, 22 2^nd^ instar reproductives, 50 3^rd^ instar reproductives and 24 pre-pupa reproductives. We then plotted the log_10_ of both measurements to see whether our classification based on morphological traits corresponded to a clear classification based on morphometric data. To further analyze the number of instars we annotated, we tested whether the growth ratios between successive instars conformed to Brooks-Dyar’s rule (Brooks 1886, Dyar 1890, Resh and Cardé 2009, Sukovata 2019). This was calculated using the following equation:

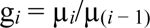

Here, g*_i_* is Brooks-Dyar’s ratio µ*_i_* is the mean head widths a given instar and µ_(*i* − 1)_ is the mean head widths for the preceding instar. Once the Brooks-Dyar’s ratio is obtained for successive instar, they are used to calculate Crosby’s growth ratio (C*_i_*), which indicates the percentage difference in growth rate between instars. This was calculated using the following equation:

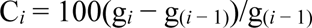

Here, difference in the change of growth exceeding ±10% between successive growth ratios indicates that the annotated instars do not conform to Brooks-Dyar’s rule. If C*_i_* is greater than 10% this indicates that one or more instars may be missing and when C*_i_* is greater than −10% this indicates that there may be too many annotated instars. Following this analysis, we collected and measured 235 additional larval samples (49 larvae from queenright colonies, 186 from queenless colonies) to see whether individuals existed between the putative 1^st^ instar and 2^nd^ worker instar, and between the 2^nd^ worker instar and the 2^nd^ reproductive instar morphospace.

### SEM microscopy

Representative instar larvae were collected from the two subcolonies used for larval collection and imaged using an environmental scanning electron microscope (FEI Inspect S SEM, Thermo-Fisher Scientific). Images were taken in a low vacuum, at an accelerating voltage of 7 kV.

### Data availability

Raw data of larval instar measurements used to generate larval instar plots is provided as Supplemental material 1.

## Results and Discussion

### Embryonic development

Under our experimental conditions (see Materials and Methods), *M. pharaonis* embryonic development lasts approximately 10 days. There are 17 embryonic stages in the fruit fly *Drosophila melanogaster* (Campos-Ortega and Hartenstein 1985). Using DAPI stained embryos (Figure 2 and 3), bright-field microscopy (Figure 4), and differential interference contrast (DIC) time-lapse videos of living embryos (Supplementary video 1), we identified homologous developmental events in *M. pharaonis*. This allowed us to annotate 17 developmental stages of *M. pharaonis* embryogenesis and harmonize them with those described in Campos-Ortega and Hartenstein (2013). These 17 developmental stages are as follows:

**Figure 2.**
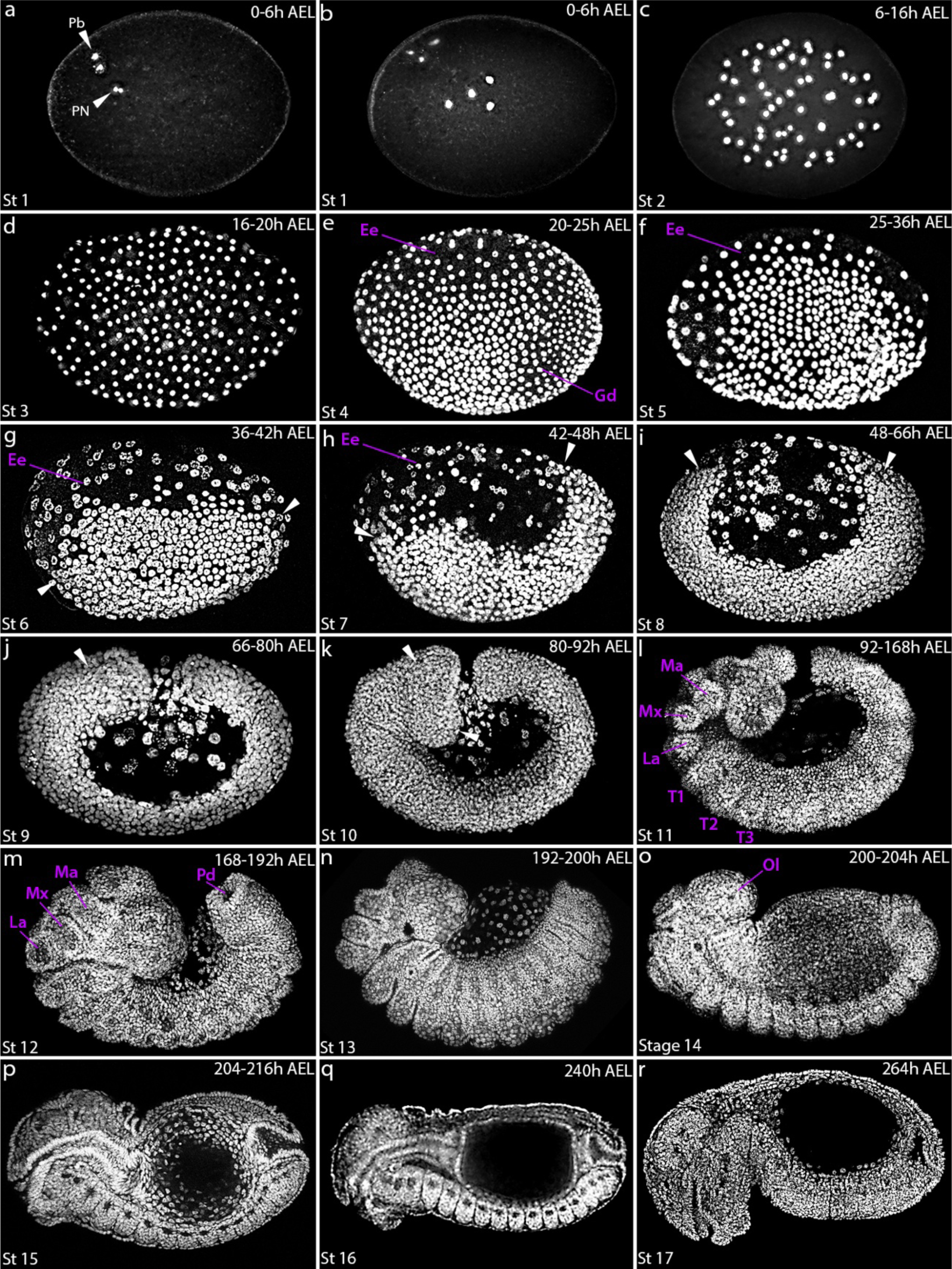
Embryonic stages of *M. pharaonis* development. a-r) Staging table of embryos stained with DAPI, which marks DNA. We harmonized *M. pharaonis* embryogenesis to the 17 stages (St) of *D. melanogaster*. Each stage is represented here. a-c) St 1– St 2 syncytial nuclear divisions. d) St 3 blastoderm stage. e,f) St 4 – St 5 cellular blastoderm stages. g,h) St 6 – St 7 gastrulation stages. i-k) St 8 – St 10 germ band extension stages. l) St 11 segmentation stage. m,n) St 12 – St 13 germ band retraction stages. o,p) St 14 – St 15 dorsal closure stages. q) St 16 embryo straightens. r) St 17, end embryogenesis Arrows in a, indicate polar bodies (Pb) and pronuclei (Pn). Arrows in g-i, indicate anterior and posterior end of the germ disc to highlight their movements toward the dorsal side of the egg. Arrows in j,k, are to highlight development of the head segment. e-h, presumptive extraembryonic region (Ee). Gd= germ disc. Mn= mandible, mx= maxilla, la= labium, pd= proctodeal invagination, T1= 1^st^ thoracic segment, T2= 2^nd^ thoracic segment, T3= 3^rd^ thoracic segment, Ol= optic lobe. Embryos are in a lateral position. Anterior is to the left, Posterior is to the right, Dorsal is up, Ventral is down. AEL = after egg laying.

**Figure 3.**
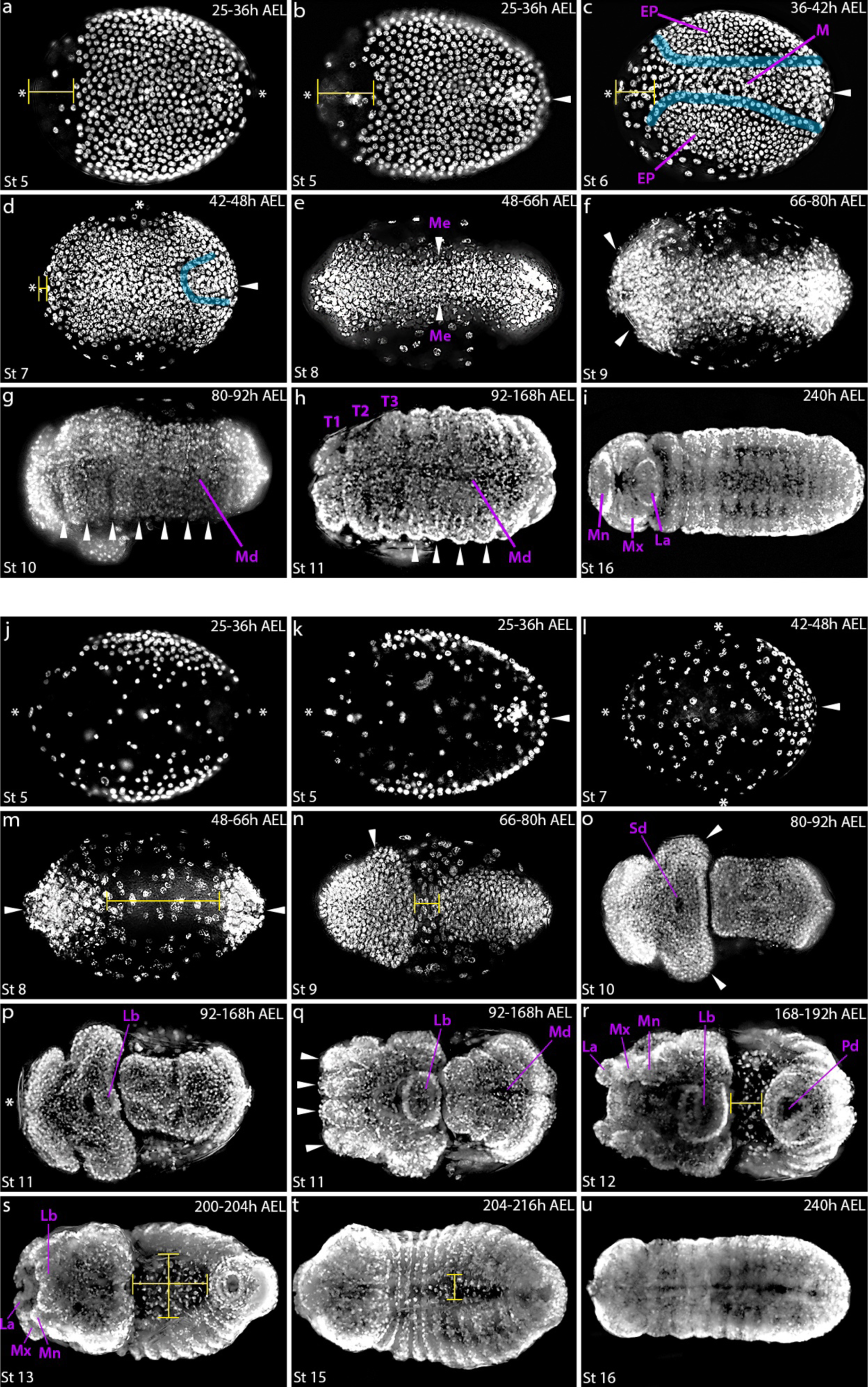
Ventral and Dorsal view of *M*. *pharaonis* germ band development. a-i) Ventral orientated embryos stained with DAPI, which marks DNA. a,b) St 5 cellular blastoderm stage embryos, asterisks highlight the absence of germ disc nuclei at the anterior or posterior pole. Arrow in b) highlights the arrival of germ disc nuclei to the posterior pole. c) St 6 Gastrulation stage. EP= presumptive ectodermal plates, M= presumptive mesoderm. Cyan line highlights the break in contact between the ectodermal plates and the mesoderm at the start of gastrulation. d) St 7 Gastrulation stage. Asterisks in the middle of the egg highlight empty space on the two lateral sides of the embryo as the ectodermal plates toward the ventral side of the embryo, which is consistent with shortening of the area marked by a cyan line that indicates the mesoderm-ectoderm boundary. a-d) Yellow brackets highlight the gradual anterior movement of the germ disc between cellular blastoderm and gastrulation stages. e) St 8 germ band extension stage. Arrows indicate the presumptive Mesectoderm (Me). f) St 9 germ band extension stage. White arrows indicate the developing head segment. g) St 10 germ band extension stage. White arrows grooves that appear to separate segment boundaries perpendicular to the ventral midline (Md). h) St 11 segmentation stage. Arrows indicate formed segments perpendicular to the ventral midline (Md). T1=1^st^ thoracic segment, T2= 2^nd^ thoracic segment, T3=3^rd^ thoracic segment. i) St 16 embryo. Gnathal segments are in the ventral direction as the embryo straightens. j-u) Dorsal orientated embryos. j) St 5 cellular blastoderm embryo. Asterisks indicates the absence of germ disc nuclei from anterior or posterior pole. k) St 5 cellular blastoderm stage. White arrow indicates movement of germ disc nuclei to the posterior pole. l) St 7 gastrulation stage. White arrow indicates a thicker layer of cells at the posterior pole, indicating that the germ disc is moving toward the posterior-dorsal side. Asterix in the middle of the egg indicate that the ectoderm has shifted toward the ventral side (compare j, k, and l). m) St 8 germ band extension stage. White arrows indicate germ disc nuclei are now at both poles. Thickness of nuclei rows indicate that the germ band is extended to the dorsal side of the egg from both poles. n) St 9 germ band extension stage. White arrow indicates development of the head anlage. o) St 10 germ band extension stage. White arrows highlight further development of the head anlage. The stomodaeum (Sd) is now visible. m,n) Horizontal yellow brackets highlight the closing of the gap between the anterior most and posterior most ends of the germ band as germ band extension progresses, until they are touching in o). p,q) St 11 segmentation stage. p) early St 11 embryo where the labrum is now visible to the posterior of the stomodaeum. q) Late St 11 embryo highlighting more developed labrum and formation of the gnathal segments (white arrows). Ventral midline (Md) is now more visible. r) St 12 germ band retraction stage. s,t) St 13 – St 15 Dorsal closure stages. Horizontal yellow bracket indicates the progression of germ band retraction between from absence in St 11 (p) to its completion in St 13 (s) embryos. Vertical yellow bracket indicates progression of dorsal closure from its absence in St 12 (r) to its completion St 15 (t). u) St 16 embryo, embryo has straightened and the gnathal segments are no longer visible. Mn= mandible, Mx= maxilla, la= labium, pd= proctodeal invagination. Anterior is to the left, Posterior to the right.

**Stage 1, syncytium.** (*0 to 6 hours after egg laying* (*HAEL*)). Freshly laid eggs are 292 ± 21 µm long and 187 ± 18 µm wide (mean ± SD, *N* = 36). The anterior pole is narrower and more rounded than the posterior pole. The four female meiotic products are located near the anterior part of the egg cytoplasm. Upon fertilization, one of the four female meiotic products (the future female pronucleus) and the male pronucleus localize together in the interior of the egg (“PN”, Figure 2a) and later fuse to form the first zygotic nucleus, similar to *D*. *melanogaster* (Kotadia et al. 2010). The other three female meiotic products (polar bodies – “Pb”, Figure 2a) remain in the cortical region of the egg (Kotadia et al. 2010). Under bright-field microscopy, the yolk appears fine and uniform in color and a small empty space is visible between the vitelline membrane and the egg cytoplasm at the posterior pole (Figure 4a). Stage 1 lasts until the end of the first two cleavage divisions, with the resulting four zygotic nuclei located in the center of the yolk (Figure 2a).

**Figure 4.**
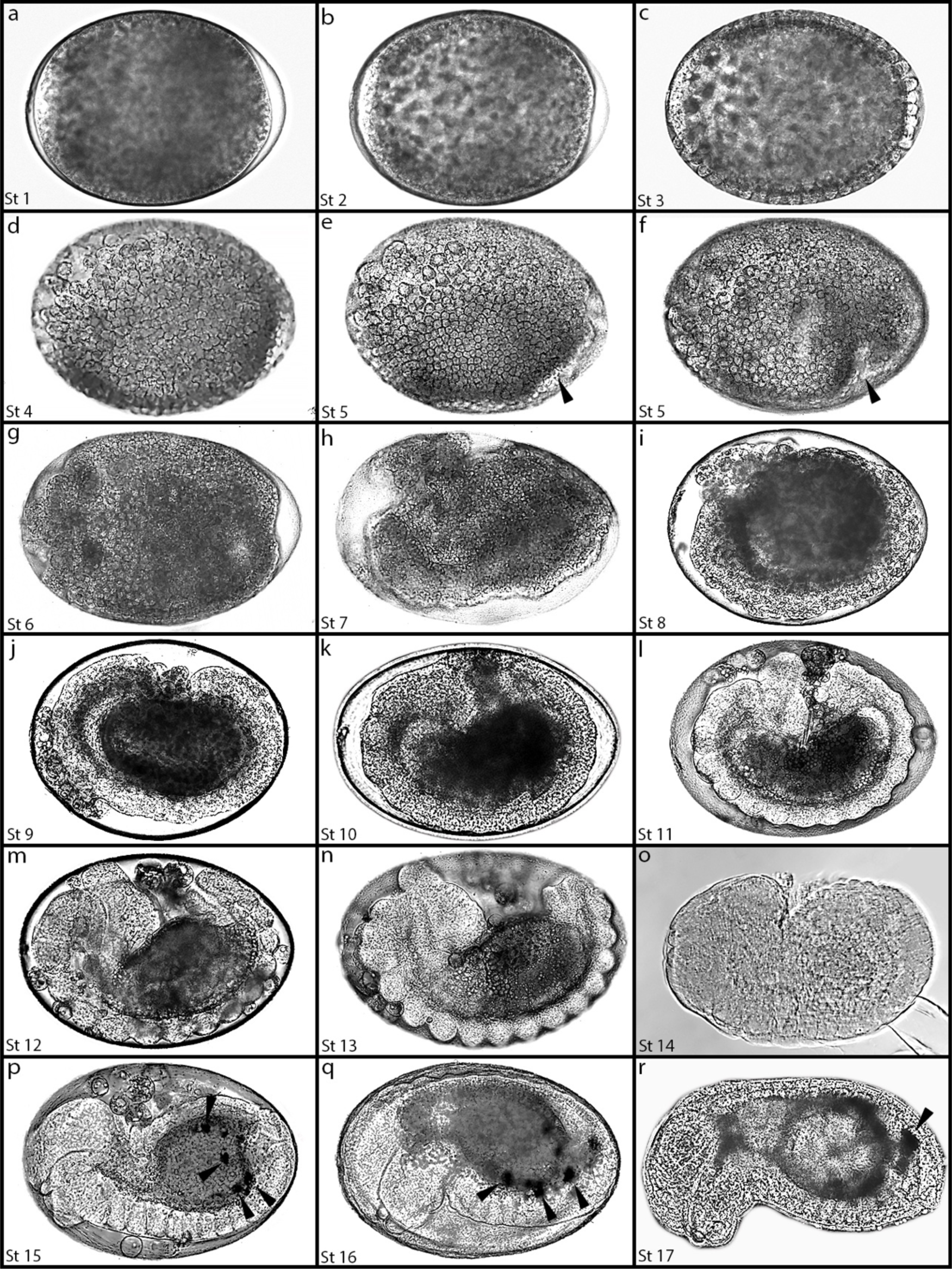
Brightfield images of *M*. *pharaonis* embryonic stages. a-r) Staging table (St) of embryos imaged using DIC. a,b) St 1 – St 2 syncytial blastoderm stages. c) Stage 3 Blastoderm stage, note change in morphology at the egg cortex. d-f) St 4 – 5 Cellular blastoderm. Note cell boundaries visible at the egg surface. Black arrow indicates formation of the ventral-posterior fold. g,f) Stage 6 – 7 Gastrulation stages. i-k) Stage 8 – Stage 10 germ band extension stages. l) Stage 11 segmentation stage. m,n) Stage 12 – Stage 13 germ band retraction stages. o,p) Stage 14 – Stage 15 dorsal closure stages. q) Stage 16. Embryo straightens. r) Stage 17. End of embryogenesis. p-r) Black arrows indicate presence of black puncta, which may be oenocytes. Anterior is to the left, Posterior is to the right, Dorsal is up, Ventral is down.

**Stage 2, syncytium.** (*6 to 16 HAEL*). In *D. melanogaster*, Stage 2 starts at the beginning of the 3^rd^ cleavage cycle and terminates at the end of the 8^th^ cleavage cycle, when the embryo consists of a syncytial blastoderm of 256 nuclei. The first synchronous nuclear divisions occur in the central part of the yolk, until 64 nuclei are formed (Figure 2c). Under bright-field microscopy, the yolk is now lighter in color during the syncytial nuclear divisions progress (compare Figure 4a to 4b). The synchronized nature of the nuclear divisions can be observed in DIC time-lapse of living embryos, as each division leads the embryo to expand and contract rhythmically (Supplementary video 1, seconds 2– 4).

**Stage 3, syncytium to syncytial blastoderm.** (*16 to 20 HAEL*). In *D. melanogaster*, Stage 3 spans the period between the last nuclear divisions and the arrival of the nuclei at the surface of the yolk (Figure 2d, Figure 4c, Supplementary video 1, seconds 5). The distribution of the nuclei at the periplasm is uniform. However, unlike in *D. melanogaster*, there is no indication of morphologically distinct pole cells at the posterior pole (Figure 2d, Figure 4c).

**Stage 4, syncytial to cellular blastoderm.** (*20 to 25 HAEL*). In *D. melanogaster*, Stage 4 is characterized by the formation of the blastoderm. During this stage, the first breaking of symmetry between the dorsal and ventral axis of the embryo occurs (Figure 2e, Supplementary video 1, seconds 6-7). The blastoderm nuclei (white) concentrate toward the ventral-lateral side of the posterior pole, whereas nuclei in the dorsal region are spaced apart. Moreover, under bright-field microscopy the cells in the dorsal-anterior region appear to be larger than those in the forming embryo primordia called the ‘germ disc’ (Figure 4d). Therefore, based on both the increased distance between nuclei and the size of cells within this region, we propose these cells to are the developing extraembryonic region of the embryo (hereafter ‘presumptive extraembryonic region’, “Ee”). However, genetic markers, such as *zen* and *dpp*, will be required to confirm the identity of these cells and determine the exact boundaries between the extraembryonic nuclei and germ disc (Panfilio 2008, Rafiqi et al. 2008). Finally, the beginning of cellularization is apparent (Figure 4d), although a membrane marker will be required to confirm whether cellularization is partial or complete.

**Stage 5, cellular blastoderm.** (*25 to 36 HAEL*). In *D. melanogaster*, Stage 5 is characterized by the completion of cellularization of the blastoderm. In *M*. *pharaonis*, further aggregation of the germ disc toward the ventral posterior side of the egg (Figure 2f), as well as an extension of presumptive extraembryonic region from the dorsal-anterior side of the egg to both the ventral-anterior and posterior pole (“Ee” in Figure 2f). Furthermore, a series of morphogenetic movements in the dorsal and postero-ventral regions can be observed (Supplementary video 1, second 8-17). First, a posterior-ventral fold forms (compare black arrows in Figure 4e to 4f and Supplementary video 1, seconds 9-11). From the dorsal view of the embryo, germ disc nuclei at the posterior pole are lacking and instead there is a presumptive extraembryonic region along the entire anterior-posterior axis (Figure 3j). Second, as Stage 5 progresses, the germ disc extends back toward the posterior pole (compare white asterix and arrow in Figure 3a to 3b and Figure 3j to 3k). Later, the dorsal-posterior region of the embryo shifts anteriorly, and the anterior side of the embryo separates from the vitelline membrane, creating a more compact embryo (Supplemental video 1, seconds 9-18).

**Stage 6, cellular blastoderm to gastrulation.** (*36 to 42 HAEL*). In *D. melanogaster*, Stage 6 encompasses gastrulation. In *M*. *pharaonis*, from the ventral view, there are apparent changes within the germ disc, where the presumptive ectodermal plates seemingly break contact with the presumptive mesoderm (“EP” and “M” in Figure 3c). The break in contact between the ectodermal plates and the presumptive mesoderm can be distinguished by two strips of more loosely organized cells between the presumptive ectoderm and mesoderm (blue line in Figure 3c). However, genetic markers such as *sog* and *twist* will be required to determine the exact boundaries between ectoderm and mesoderm (Stathopoulos and Newcomb 2020). Furthermore, during this stage the space between the anterior pole of the egg and the anterior of the germ disc decreases, as the anterior of the germ disc extends antero-dorsally (compare yellow brackets in Figure 3a-c and to Figure 4g). In the lateral position, the germ disc becomes more organized forming a wedge that takes up about 2/3 of the egg, while the presumptive extraembryonic region makes up the rest of the egg, including the entire dorsal side (Figure 2g). Finally, the embryo becomes more distant from the anterior of the vitelline membrane, where it becomes even more compact in shape (Supplementary video 1, 19 to 29 seconds).

**Stage 7, gastrulation to germ band extension.** (*42 to 48 HAEL*). In *D. melanogaster*, Stage 7 is characterized by the end of gastrulation and initiation of germ band elongation. *M. pharoanis* also undergoes both of these processes at this stage, but unlike *D. melanogaster*, gastrulation proceeds with no signs of mesoderm invagination. Instead, it appears as though the ectodermal plates slide over the mesoderm resulting in its internalization (Compare cyan line in Figure 3c to 3d). This is further evidenced by looking at both ventral and dorsal views, where germ disc nuclei are absent from the lateral sides of the egg (compare Figure 3b to 3d and Figure 3k to 3l; asterisks denotes absence of presumptive ectoderm). This suggests that the presumptive lateral ectoderm has shifted ventrally. During this stage, germ band elongation initiates, where both anterior and posterior ends of the germ disc extend along the dorsal side of the egg (compare white arrows in Figure 2g to 2h, Figure 4g to 4h, and Supplemental video 1, 30 to 37 seconds). As a result, the presumptive extraembryonic region is shifted toward the antero-dorsal compartment of the egg (Figure 2h). From the ventral view, as the germ band extends toward the anterior pole of the egg, the gap between the embryo and the anterior of the egg becomes almost completely closed (compare yellow brackets in Figure 3c to d).

Gastrulation itself in *M. pharaonis* appears to follow the “hymenopteran type” reported for honeybees (Fleig and Sander 1986; Fleig and Sander 1988) and described by Lynch in *Nasonia vitripennis* (Lynch et al. 2012). For example, the blastoderm cells located in the dorsal and lateral sides move in a ventral direction, forming a compact epithelium covering the ventro-lateral regions of the blastoderm (compare Figure 3b-d). This movement leaves very few blastoderm nuclei at the dorsal region (Figure 3l). The morphology of the dorsal region appear to be similar to the “dorsal strip” characterized in *A. mellifera* (Fleig and Sander 1986).

**Stage 8, germ band extension.** (*48 to 66 HAEL*). In *D. melanogaster*, Stage 8 is characterized by the continuation of germ band elongation. In *M. pharaonis*, the extension of the germ band toward the dorsal side of the egg continues (Figure 2i, Figure 4i, Supplemental video 1, 38 seconds to 1 minute 30 seconds). As the anterior end of the germ band shifts dorsally, the posterior end of the germ band shifts slightly ventrally from its maximal dorsal position in Stage 7, resulting in a symmetrical embryo (compare white arrows in Figure 2h to 2i). Unlike in Stage 7, where only the posterior end of the germ band is at the dorsal side of the egg, in Stage 8, both anterior and dorsal ends of the germ band are dorsal (compare asterisks and white arrows in Figure 3l to 3m). On the ventral side the germ band narrows, presumably because the ectoderm plates have now completely slide over the mesoderm following gastrulation in stage 7 (Figure 3e). Furthermore, at the surface of the ventral view there is a narrow overlay of ectoderm ventral to the mesoderm (white arrows in Figure 3e). We infer this to be the presumptive mesectoderm. However, this would require confirmation with a suitable genetic marker, such as expression of the *single-minded* gene (Stathopoulos and Newcomb 2020).

**Stage 9, germ band extension.** (*66 to 80 HAEL*). In *D. melanogaster*, Stage 9 is characterized by the completion of germ band extension. In *M*. *pharaonis*, the germ band has reached its maximum extension and it now covers the dorsal side of the egg (Figure 2j and Figure 4j, Supplemental video 1 1 minute 31 seconds to 1 minute 39 seconds). From the dorsal view, the gap between the anterior and posterior ends of the germ band is almost closed (compare yellow brackets Figure 3m to 3n). Furthermore, both ventral and dorsal views reveal that the head anlage is beginning to develop at the anterior of the germ band, where the anterior end becomes markedly thicker than the posterior of the germ band (white arrow in Figure 3f and Figure 3n).

**Stage 10, segmentation.** (*80 to 92 HAEL*). In *M. pharaonis*, Stage 10 is characterized by continued development of the germ band (Figure 2k and Figure 4k). From the dorsal view, the anterior and posterior ends of the germ band are now in contact (Figure 3o). Both lateral and dorsal views show a significant increase in the size of the head anlage (compare white arrows in Figure 2j to 2k and Figure 3n to 3o). Moreover, from the ventral view we observe a series of grooves running perpendicularly to the embryonic midline, creating boundaries between germ band segments (white arrows in Figure 3g). Finally, the stomodaeum (precursor of the mouth) arises as an ovoidal invagination (“sd”, Figure 3o).

**Stage 11, segmentation.** (*92 to 168 HAEL*). In *D. melanogaster*, Stage 11 is characterized by the completion of segmentation, where all segments of the embryos become well defined. In the lateral view, the gnathal segments in *M*. *pharaonis* (mandible “mn”, maxilla “mx”, and labium “la”) are well defined as rounded bulges (Figure 2l), while the thoracic and abdominal segments are less pronounced. The thoracic segments (T1, T2, T3) are be more easily discernible than the abdominal segments (Figure 2l). However, by viewing the embryo from the ventral side, the pronounced ridges between abdominal segments are now discernible (arrows in Figure 3h). Furthermore, bright-field microscopy shows even clearer segmentation through the entire embryonic abdomen (Figure 4l). Moreover, the ventral midline is now visible (“md” in Figure 3h). From the dorsal view, the head anlage develops more pronounced folds (Figure 3q). Specifically, the stomodaeum deepens, and the labrum begins to form as an ovoid protrusion of tissue immediately anterior to the stomodaeum (“lb” in Figure 3p). As stage 11 progresses, the gnathal segments continue to develop (compare the asterisk in Figure 3p to arrows in Figure 3q).

**Stage 12, germ band retraction.** (*168 to 192 HAEL*). In *D. melanogaster*, Stage 12 is characterized by the start of germ band retraction. In *M. pharaonis*, the distance between the posterior end of the germ band from the posterior end of the head anlage increases (Figure 2m, horizontal yellow bracket, Figure 3r, Figure 4m). During this stage, the gnathal segments appear elongated, are oriented upward, and are condensed (Figure 2m, Figure 3r). Furthermore, the labrum grows significantly (Figure 3q). Moreover, abdominal segments are now easily delineated when viewing the embryo laterally (Figure 2m). Finally, the proctodaeum becomes visible (“Pd” Figure 2m, Figure 3r Figure 4m).

**Stage 13, germ band retraction.** (*192 to 200 HAEL*). In *D. melanogaster*, Stage 13 is characterized by the end of germ band retraction and the start of dorsal closure. In *M. pharaonis*, there is a further increase in the gap between the posterior end of the germ band from the posterior end of the head anlage (Figure 2n, compare horizontal yellow brackets Figure 3r and 3s and Figure 4n). Conversely, at the onset of dorsal closure, there is a narrowing of the two lateral flanks of the embryo as they grow toward the midline (compare vertical yellow brackets in Figure 3r to 3s). Finally, the gnathal segments and labrum orient toward the anterio-ventral side of the embryo (Figure 2n, compare Figure 3r to 3s).

**Stage 14, dorsal closure.** (*200 to 204 HAEL*). During Stage 14, the gnathal segments are now oriented toward the anterior of the egg, as the head shifts towards the ventral side (Figure 2o, Figure 4o). Optic lobe (“Ol”) is now visible in the head (Figure 2o). Dorsal closure proceeds (Figure 2o).

**Stage 15, dorsal closure.** (*204 to 216 HAEL*). In *D. melanogaster*, Stage 15 is characterized by the completion of dorsal closure. In *M. pharaonis*, the lateral flanks meet at the dorsal midline to close the dorsal side of the embryo (Figure 3t). Unlike *Drosophila*, which undergoes head involution, the gnathal segments continue to orient ventrally, and now face toward the ventral side in *M. pharaonis* embryos (Figure 2p, Figure 4p). Under bright-field microscopy, small black spots are present in the posterior region of the gut (black arrows in Figure 4p). Although a genetic marker would be required to determine the nature of these black spots, based on their morphology and localization, we propose that they are oenocytes, which are black spots observable by bright-field miscroscopy in the posterior abdominal segments of larval instars (data not shown).

**Stage 16, straightening.** (*10 days after egg laying*). During stage 16, the embryo straightens (Figure 2q, Figure 3i, Figure 3u, Figure 4q) and the gnathal segments are oriented ventrally (Figure 3i). The black dots persist in the posterior of the gut (black arrows Figure 4q).

**Stage 17, micro larvae.** (*11 days after egg laying*). Stage 17 marks the end of embryogenesis. The fully formed 1^st^ instar larvae outgrow the chorion and hatch (Figure 2r, Figure 4r). The black dots coalesce toward the dorsal posterior of the embryo (black arrow in Figure 4r).

### Germ cell specification

Across insects, germ cells are specified via one of two major modes: 1) preformation, where a specialized cytoplasm called germplasm is maternally deposited into developing oocytes to specify germ cells, or, 2) induction, where cell-to-cell signaling induces germ cell identity via zygotic mechanism later in embryogenesis (Extavour and Akam 2003). Overall, the macroevolutionary patterns of how different insect lineages specifies their germline continues to be elucidated. Within Hymenoptera, while both modes are present, preformation appears to be more common (Khila and Abouheif 2008, 2010, Lynch et al. 2011, Rafiqi et al. 2020). To characterize how germ cells are specified in *M*. *pharaonis*, we used *in situ hybridization* to first detect *nanos* mRNA, which marks the germline across animals (Extavour and Akam 2003). In early embryos (Stage 1 and 2), similar to other ants described to date we find *nanos* localized to the posterior pole marking germplasm, similar to other ants described to date (Figure 5a and b). Surprisingly, during Stage 3 and 4, when germ cells form extruded pole buds in *D*. *Melanogaster*, *M*. *pharaonis* germ cells remain at the posterior cortex of the embryo and do not bud off (Figure 5c and d). Instead, during the cellular blastoderm stage (Stage 5), we find two alternative patterns of germ cell localization in stage-matched embryos (Figure 5e-f’): one where germ cells are located inside the embryo, which we call ‘in-phenotype’ embryos (n=62; Figure 5e and e’) and one where germ cells protrude from the posterior pole, which we call ‘out-phenotype’ (n=124; Figure 5f and f’). Next, to confirm the identify of these cell clusters as bonafide germ cells, we used two additional germline markers, *oskar* mRNA and Vasa protein, both of which strongly mark germ cells either inside or outside the embryo (Figure 5g-h’). Finally, to rule out the possibility that the in-phenotype and out-phenotype embryos are not successive developmental stages (i.e. in-to-out or out-to-in), we used live imaging to see if we can morphologically distinguish different germ cell migration patterns between embryos (Figure 5i-j’’’ and Supplementary video 2 and 3). Live imaging confirms that in-phenotype and out-phenotype embryos are two alternative phenotypes and not successive stages. For in-phenotype embryos, we find no extruded cells are detected at the posterior pole, instead we observe clearing inside the cellular blastoderm consistent with the internal location of *nanos*-marked germ cells (Figure 5f, black arrows in Figure 5i-I’’’, Supplementary video 2). Four out-phenotype embryos, we find cells that are extruded at the posterior pole consistent with the external location of *nanos*-marked germ cells (Figure 5f’, white arrows in Figure 5j-j’’’, Supplementary video 3). Taken together, our results reveal the discovery of a germ cell migration polyphenism in *M*. *pharaonis* embryos.

**Figure 5.**
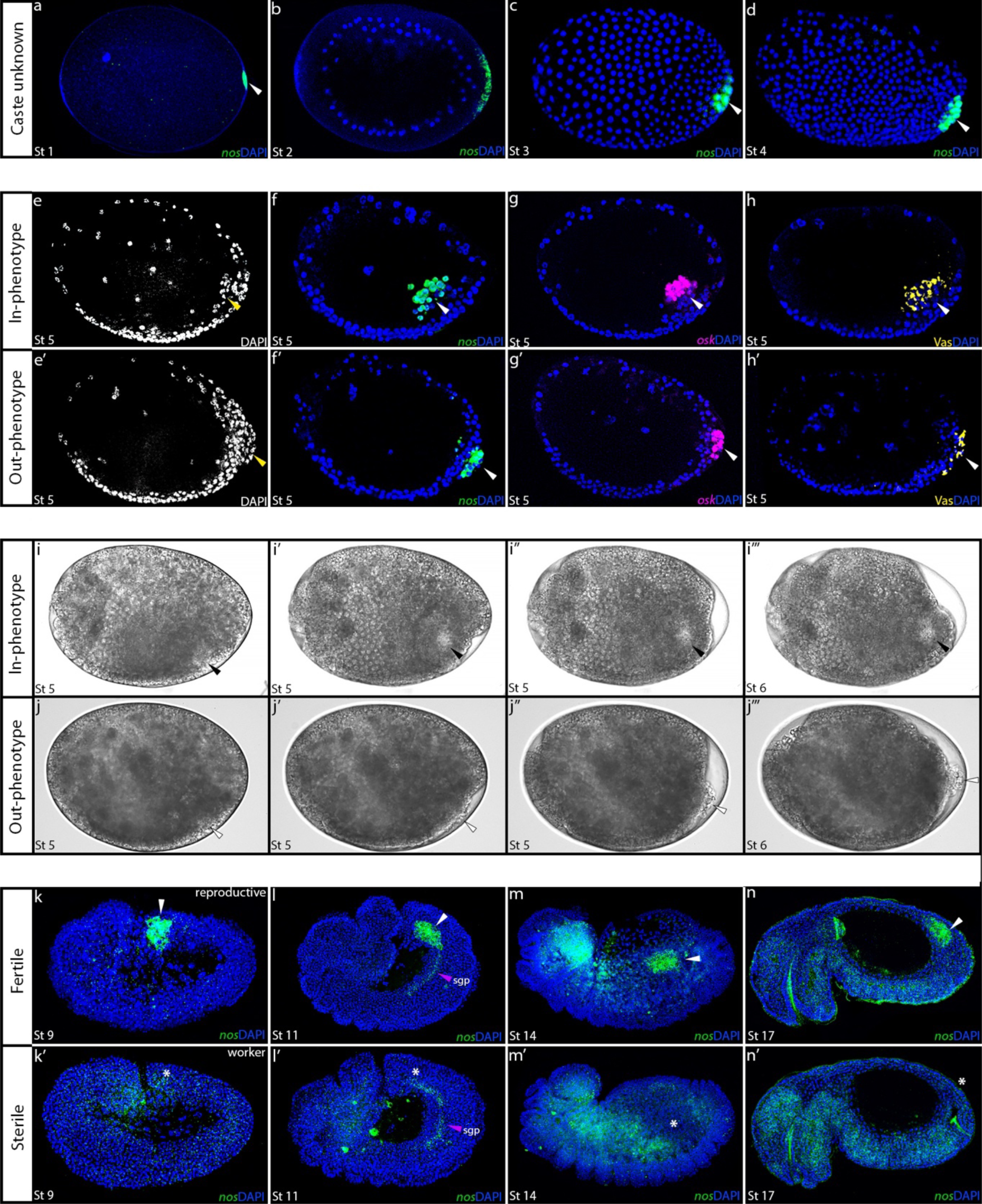
A Germ cell migration polyphenism in underlies embryonic caste differentiation in *M*. *pharaonis*. a-d) Early *M*. *pharaonis* embryos stained DAPI (blue) and *nanos* (*nos*, green) to reveal germ cell specification. a) St 1 syncytial embryo showing *nos* mRNA in germplasm at the posterior cortex. b) St 2 syncytial embryo showing nuclei approaching germplasm (*nos* mRNA) at the posterior cortex. c) St 3 embryo showing initial formation of germ cells (*nos* mRNA). d) St 4 embryo showing fully specified germ cells at the posterior of the developing germ disc. e-h’) Germ cell polyphenism during cellular blastoderm stage (St 5). ‘In’-phenotype (e, f, g, h) and ‘Out’-phenotype (e’, f’, g’, h’) embryos stained with DAPI (e, e’), *nos* mRNA (f, f’), *oskar* (*osk*) mRNA (g, g’) or Vasa (Vas) protein (i, i’). i-j’’’) Live-imaging of ‘In’ and ‘Out’-phenotype embryos. i-i’’’) Successive frame captures of an ‘In’-phenotype embryo. Black arrow highlight clearing that coincides with the germ cell cluster. j-j’’’) Successive frame captures of an ‘Out’-phenotype embryo. White arrow highlights cluster of cells extruded for the posterior of the embryo that coincide with the germ cell cluster. Both embryos start at Stage 5 cellular blastoderm stage and end at Stage 6 gastrulation stage. See supplementary video 2 to 3 for respective videos. k-n’) Developmental trajectory of germ cells stained with *nos* mRNA in post-gastrulated embryos. k, l, m, n) Fertile embryos with *nos* mRNA marked germ cells (white arrow). k’, l’, m’, n’) Sterile embryos that lack germ cells (Asterix *), however *nos* mRNA continues to mark the nervous system and the somatic gonadal precursor cells (sgp, magenta arrow). Anterior is to the left, Posterior is to the right, Dorsal is up, Ventral is down.

*M*. *pharaonis* is a species with an obligate sterile worker caste, meaning that workers completely lack a germline and do not develop ovaries. Social evolutionary theory predicts that the larger the queen-worker asymmetry within a given species, the earlier in developmental time caste determination and differentiation occurs (Wheeler 1986). Consistent with this theory germ cells could not be detected in embryos from another *Monomorium* species, *M. emersoni* (Khila and Abouheif 2010). Our discovery of two unique germ cell localization patterns raises the exciting possibility that a germ cell migration polyphenism may underly caste determination during early embryogenesis to give rise to caste-differentiated larvae. To test this hypothesis, we probed mid-to-late-stage embryos (Stage 8 to 17) for *nanos* mRNA to determine whether we could identify fertile (embryos with germ cells) or sterile (embryos without germ cells) individuals (Figure 5k-n’). From the germ band extension stage (Stage 8) onwards we found two populations of stage-matched embryos with and without germ cells, showing the presence of fertile and sterile embryos that we interpret as reproductives (gyne or male) and workers, respectively (Figure 5k-n’). Taken together, the finding of both two locations of germ cells during the blastoderm stage and fertile or sterile embryos later in embryonic development strongly suggest that localization of germ cells within the embryo is one of, if not the, earliest point in caste differentiation between reproductives and workers.

Overall, our findings show that with respect to reproduction, gyne/queen and worker castes are already differentiated during early embryogenesis, however, it is difficult to ascertain from our current knowledge of model organisms whether the in-phenotype or out-phenotype represents the queen or worker state. Within holometabolous insects germ cells can either from budded off from the blastoderm, as is the case for *D*. *melanogaster* ‘pole cells’, or within the posterior of the blastoderm, as is the case for *Tribolium castaneum* (Schröder 2006, Stappert et al. 2016, Benton et al. 2019, Campos-Ortega and Hartenstein 2013; Figure 6c-f). Therefore, further characterizing the developmental fates of in-and-out germ cells will be required in future studies.

**Figure 6.**
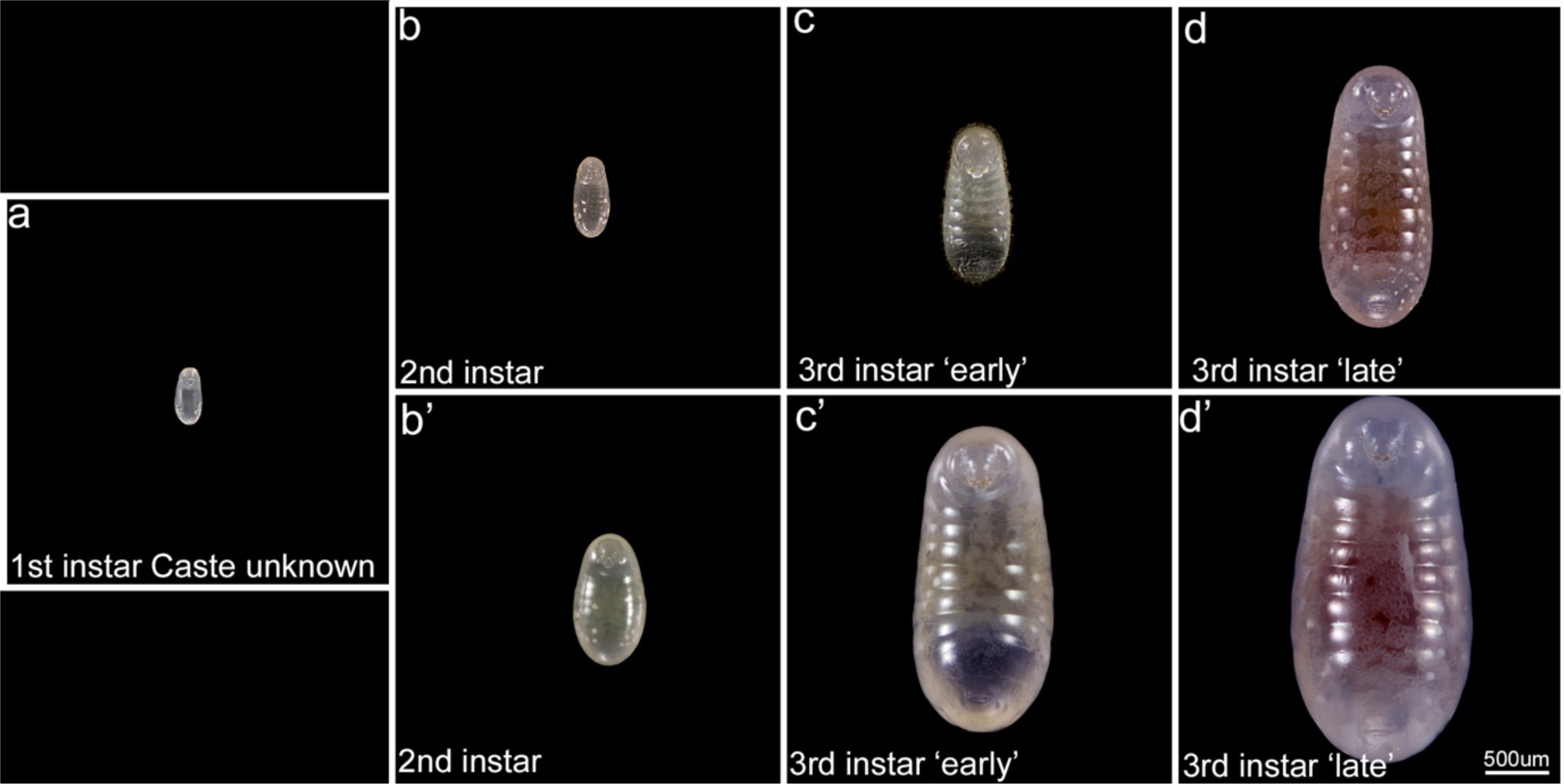
Brightfield images of *M*. *pharaonis* larval instars. Instar designation is based on Berndt and Eichler 1987, Alvares et al. 1993. a) 1^st^ instar larvae ‘Caste unknown’, where caste cannot be determined based on morphological characters. b-d) worker-destined larvae. b) 2^nd^ instar larvae. c) 3^rd^ instar larvae ‘early’. d) 3^rd^ instar larvae ‘late’. B’-d’) reproductive-destined larvae. f) 2^nd^ instar larvae. g) 3^rd^ instar larvae ‘early’. h) 3^rd^ instar larvae ‘late’. All images are to scale.

### Larval development

Under our experimental conditions larval development in *M*. *pharaonis* lasts approximately 22 days. Previous studies have provided information regarding the number and morphology of larval instars in *M. pharaonis*, making use of both morphometric measurements (i.e. head width, body length, diameter of the first thoracic spiracle) and examination of types and number of cuticular features such as setae, spines and tuberculi (Berndt and Eichler 1987, Alvares et al. 1993). The combined use of these traits is often sufficient to determine the number of larval instars in ants (Masuko 2017). On the basis of these studies, *M. pharaonis* has been characterized as having three larval instars (Alvares et al. 1993, Figure 6). Berndt and Eichler (1987) provide thorough descriptions of *M*. *pharaonis* larvae. However, these descriptions are only available in German, therefore, we provide brief characterizations of worker and sexual larvae based on his descriptions and the examination of SEM images (Figure 7).

**Figure 7.**
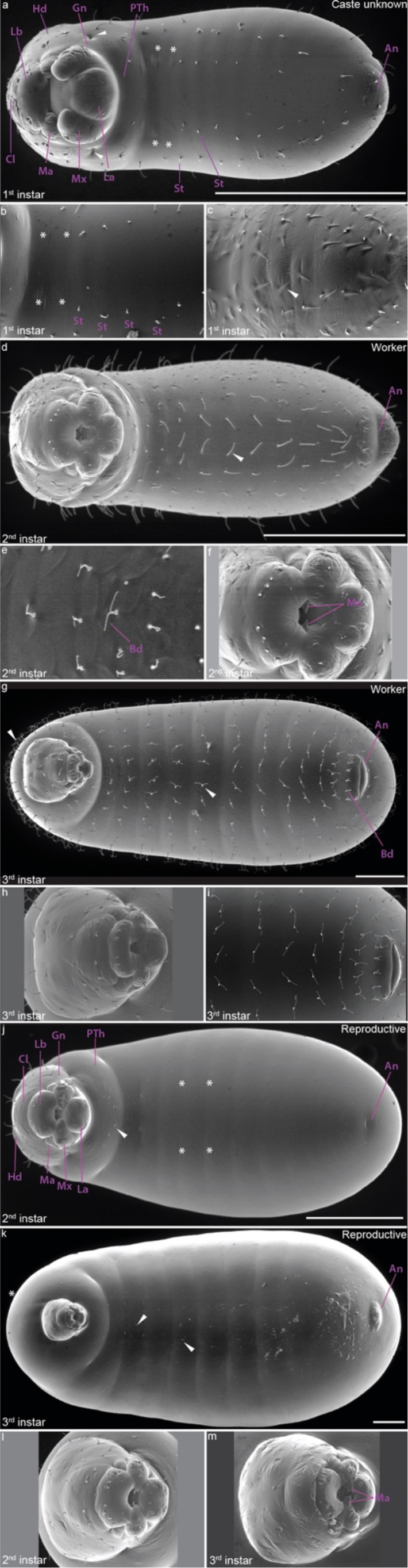
SEM images of *M*. *pharaonis* larval instars. Instar designation is taken from Berndt and Eichler 1987. a-c) 1^st^ instar larvae. a) 1^st^ instar larvae. b) Zoom in of thoracic region of early first instar larvae. c) Zoom in of thoracic region of late first instar larvae. d-f) 2^nd^ instar worker larvae. d) 2^nd^ instar worker larvae. e) Zoom in of thoracic region of late 2^nd^ instar worker larvae. f) Zoom in of mouthparts of early 2^nd^ instar worker larvae. g-i) 3^rd^ instar worker larvae. g) 3^rd^ instar worker larvae. h) Zoom in of head and mouth parts of 3^rd^ instar worker larvae. i) Zoom in of abdominal region of 3^rd^ instar worker larvae. j) 2^nd^ instar reproductive larvae. k) 3^rd^ instar reproductive larvae. l) Zoom in of head and mouthparts of 2^nd^ instar reproductive larvae. m) Zoom in of head and mouthparts of 2^nd^ instar reproductive larvae. White arrow in a) indicates slit-like opening at border between gena and prothorax. Asterisks indicates absence of hairs on the inside of the 2^nd^ and 3^rd^ thoracic segments. White arrow in c) indicates subcuticle hair that will appear after molting. White arrow in d) indicates long simple hair. White arrow in g) indicates anchor hair. White arrow in j, k) indicate rare hairs on reproductive larvae. Hd= head, An= anus, St= setae, Gn= gena, PTh= prothorax, Bd= bifid, Ma= mandible, Mx= maxilla, La= labium, Cl= clypeus, Lb= labrum. White line indicates scale of 100um.

**First instar larvae (caste not discernible)**: 1^st^ instar larvae are slightly longer on the major axis than a freshly laid egg (length ± SD: 0.389 ± 0.032 mm), with an average head width of 0.142 ± 0.006 mm (n = 103). They are whitish in pigmentation and fat with a broader posterior than putative 2^nd^ instar workers (compare Figure 6a and b). The head is ventral at the anterior end, while the anus is postero-ventral (head “Hd”, anus “An” Figure 7a). They have very few cuticular body hairs that are sparse and simple in morphology (Figure 7b). On the ventral side, the prothorax and the abdominal segments bear four short setae, roughly organized in two rows across the ventral midline (setae “St” in Figure 10a and b). The second and third thoracic segments always lack the short hair in the inner row (asterisks in Figure 7a and b), which is a distinct trait of this instar. The head is simple, with a slit-like opening at the border between the gena (the lower part of the head that extends behind the maxilla) and the prothorax (white arrows, gena “Gn”, and prothorax “PTh” in Figure 7a). Note that as the first instar is ready to molt, we observe an increase in the density of hairs and length of setae along the ventral side of the larvae (white arrow, Figure 7c).

**Second instar larvae (worker and reproductive destined)**: 2^nd^ instar worker-destined larvae have an average length of 0.572 ± 0.090 mm and an average head width of 0.173 ± 0.008 mm (N=51). They are whitish in pigmentation and more slender than a 1^st^ instar at the posterior end (Figure 6b, compare Figure 7a to 7d). The location of the anus is sub-terminal. We observed numerous long and simple body hairs, rarely bifid (y-shaped), that are mostly uniformly distributed and organized in rows along the segmented body (white arrow, compare ventral surface in Figure 7a to d, as well as to bifid “Bd” in Figure 7e). Unlike in the 1^st^ instars, hairs are present on the midline of the ventral surface (compare Figure 7a to b, 7d to e). We observed few simple and sparse hairs on the head. Compared to the 1^st^ instar, the mandible tooth has become more pointed (“Ma” in Figure 7f) Second instar reproductive-destined larvae have an average length of 1.026 ± 0.192 mm and head width of 0.215 ± 0.004 mm (n=22). They are whitish in pigmentation and larger at the posterior end compared to a 2^nd^ instar worker-destined larvae (compare Figure 6b and b’, and compare Figure 7d to j). Unlike 2^nd^ instar worker-destined larvae, however, we found no hairs on thoracic and abdominal segments, with the exception of the prothorax that bears few short, simple hairs on the ventral side (white arrow and asterisks in Figure 7j). Moreover, we observed few simple and sparse hairs on the head, organized in a similar fashion to 2^nd^ instar worker-destined larvae.

**Third instar (worker and reproductive destined)**: 3^rd^ instar worker-destined larvae have an average body length of 1.551 ± 0.432 mm and an average head width of 0.273 ± 0.009 mm (n=55). They are whitish in pigmentation, but unlike previous instars the gut pigmentation is becoming increasingly darker, and mandibles are now darker (Figure 6d). We observed numerous bifurcated, anchor-shaped hairs on the cranium, thoracic and abdominal segments, organized in rows following body segmentation similar to the simple hairs 2^nd^ instar worker-destined larvae (compare white arrows in Figure 7d and g). Moreover, we observed bifid hairs around the anus (Figure 7g and i). Finally, the clypeus and mouth parts are similar in terms of shape to those described in 2^nd^ instar worker-destined larvae (compare Figure 7f to h). Third instar reproductive-destined larvae have an average body length of 2.464 ± 0.397 mm and an average head width of 0.284 ± 0.014 mm (n = 50). They are whitish in pigmentation and similar to 3^rd^ instar worker-destined larvae with gut pigmentation that becomes increasingly darker with age and pigmentation of the mandibles (Figure 7k). Similar to 2^nd^ instar reproductive-destined larvae, they are more rotund than 3^rd^ instar worker-destined larvae. We observed extremely short, straight hairs on thoracic and abdominal segments, that can only be detected using SEM microscopy (Figure 6d’ and white arrows in Figure 7k). The cranium possesses almost no hairs compared to 2^nd^ instar reproductive larvae (asterisks in Figure 7k). Finally, the mandibles become serrated and increasingly darken with age (Figure 6d’ and 7m).

Next, using morphological criteria described in Berndt and Eichler (1987) to sample larvae, we characterized the larval instars of *M*. *pharaonis* worker and reproductives by plotting measurements of the maximum head width versus the maximum larval length along the antero-posterior axis of 340 larvae (Figure 8a). Consistent with Berndt and Eichler (1987) and Alvares et al. (1993), we found 3 larval instars (Figure 8a). While larvae of *M*. *pharaonis* having 3 instars is consistent with other *Monomorium* species, such as *M. floricola* and *M. trivale* (Berndt and Eichler 1987, Alvares et al. 1993, Solis et al. 2010, Idogawa et al. 2022), we identified discrepancies regarding the growth rates between instars in both workers and reproductive larvae. Specifically, we found that the inter-instar growth rates are counter intuitive between worker and reproductive larvae. For example, while the growth between 1^st^ instar larvae and 2^nd^ instar reproductive-destined larvae is more than double that of 2^nd^ instar worker larvae (72.43µm vs 32.1µm), the growth between the 2^nd^ and 3^rd^ instar is 25% greater for workers-destined larvae than reproductives (93.03µm vs 69.98µm). The possibility that worker-destined larvae grow 25% more between penultimate and final instars than reproductive-destined larvae is counterintuitive to the metabolic needs required to generate the larger body sizes of gynes. To some extent, this discrepancy in instar annotation may be a unique problem to *M. pharaonis* because larvae of all three castes (i.e. workers, gynes, and males) are produced simultaneously and constantly. As such, there may be instances where differentiating between castes is difficult.

**Figure 8.**
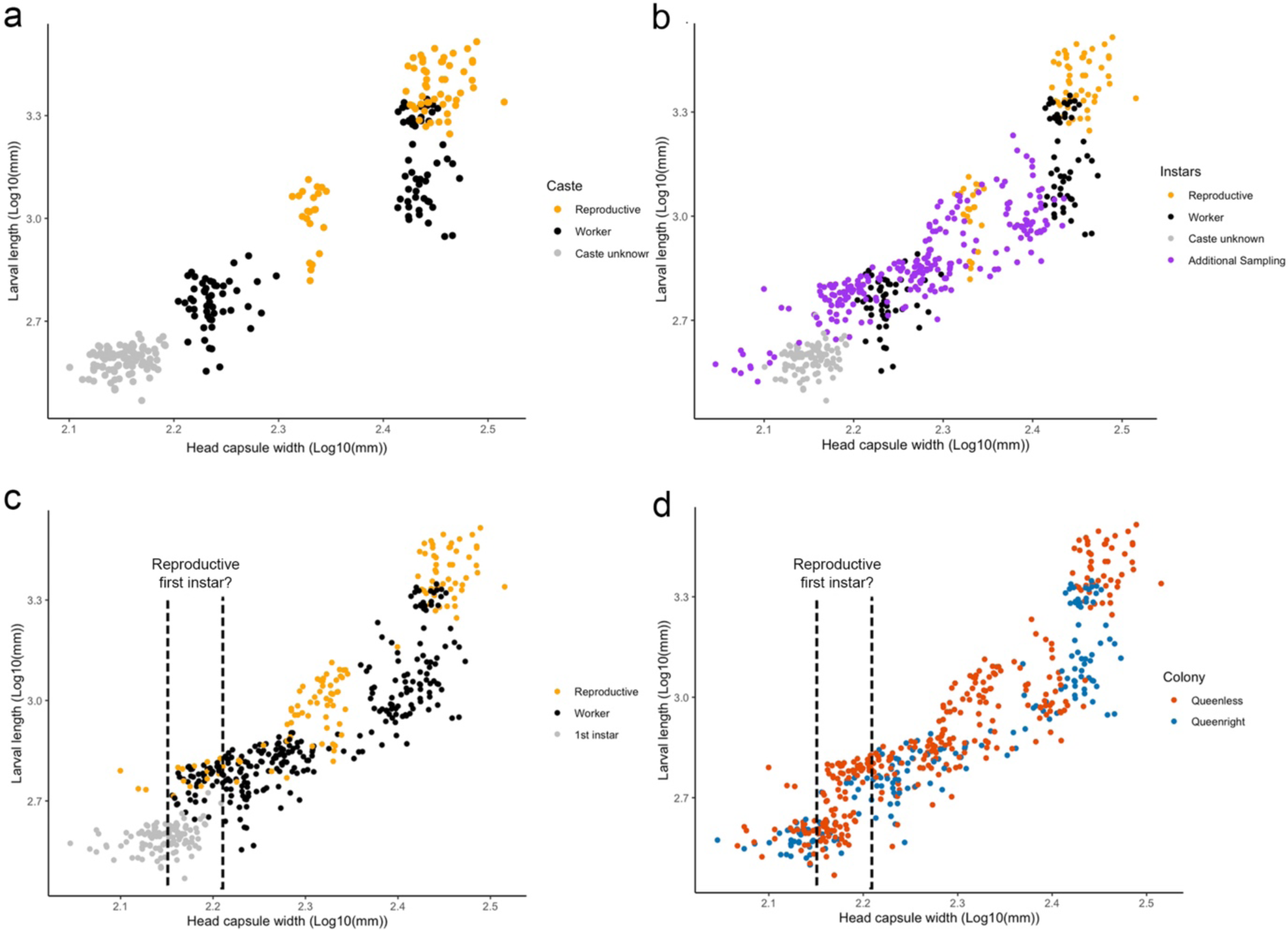
*M. pharaonis* larval instar analysis. a) Plot of log Larval length by log Head capsule width to visualize instar clusters of 340 larvae, sampled according to Berndt and Eichler 1987. White = Caste unknown, gold= reproductive-destined caste, black = worker-destined caste. This plot shows 3 instars for each caste, where the 1^st^ instar is morphologically ambiguous. b) Plot of log Larval length by log Head capsule width to visualize instar clusters with 235 additionally sampled larvae (Additional sampling, purple). c) Plot of log Larval length by log Head capsule of all 575 sampled larvae, annotated by caste according to their morphological landmarks. d) Plot of log Larval length by log Head capsule of all 575 sampled larvae, annotated by colony-type they were sampled from. Red = queenless, Blue = queenright. Vertical dotted line indicates proposed 1^st^ reproductive-destined larvae morphospace.

To better understand whether our annotation of *M*. *pharaonis* instars was incomplete, we tested whether changes in mean head width adhered to the Brooks-Dyar’s rule (Brooks 1886, Dyar 1890, O’Neal and Markin 1975). There are two components to the Brooks-Dyar’s rule. The first is to statistically test whether growth between instars is linear, such that when the natural logarithm (Ln) of mean head width of each instar is plotted using linear regression, R^2^ should be close to 1. We found that the model is linear with a good fit of R^2^ = 0.9983 for worker-destined and R^2^ = 0.9954 for reproductive-destined larvae. The second component of Brooks-Dyar’s rule states that any highly sclerotized structure, such as the head capsule, increases between each instar by a constant growth ratio (Brooks 1886, Dyar 1890, Resh and Cardé 2009). Deviations from a constant growth ratio can be detected by calculating the Brooks-Dyar’s ratio and then using this ratio to calculate Crosby’s growth ratio, where deviations > 10% indicates that an instar may be missing, while a deviation of >-10% indicates there may be too many annotated instars (Sukovata 2019). We found a difference that exceeds ±10% between successive growth ratios for both worker and reproductive-destined larvae, indicating a failure to adhere to Brooks-Dyar’s rule. For the worker-destined larvae we calculated a Crosby’s growth ratio of 25.14%, and for reproductive-destined larvae we calculated a growth ratio of −12.16%. Together, these results suggest that based on morphological landmarks alone our instar identification is incomplete.

In order to determine whether we excluded important caste ambiguous larvae by relying on morphological landmarks alone, we sampled an additional 235 larvae irrespective of morphological landmarks (e.g. rounded, presence of hairs) and measured maximum head width and larval length (Figure 8b and c). We found that additional sampling filled the gaps in head width morphospace between our previously annotated instar clusters. We then repeated our calculations for the Crosby’s growth ratios to include our additional larvae. We found that for worker-destined larvae the ratio only slightly improved from 25.14% to 20.6 percent. This result suggests that more precise metrics, such as ecdysone titres may be required to determine the worker-destined larvae instar boundaries. On the other hand, for reproductive-destined larvae we found that our additional sampling substantially improved the Crosby’s growth ratio from −12.16% to 7.99%, where now reproductive-destined larvae growth adheres to Brooks-Dyar’s rule.

Finally, our additional sampling revealed a cluster of larvae that overlap in head capsule size with caste unknown 1st instar larvae but possess larval lengths similar to our previously annotated 2^nd^ instar worker-destined larvae (see dotted lines Figure 8c). Surprisingly, within this cluster a handful of larvae can be further annotated as reproductive-destined based on their morphology. When we code our samples based the colony-type they were collected from (queenless or queenright) we find that this cluster is almost entirely composed of larvae collected from queenless colonies (Figure 8d). This is significant as queenless conditions increase the likelihood for developing larvae to be reproductive, as they are normally culled under queenright conditions. Taken together, these results raise the possibility that within the 1^st^ instar, putative reproductive-destined larvae accelerate their growth to establish a reproductive-specific growth trajectory. Given that we identified that reproductive and worker castes are differentiated during embryogenesis, subsequent growth within the 1st larval instar may be necessary to realize a strong queen-worker body size asymmetry.

The nearly continuous distribution in maximum head capsule size of both worker and reproductive-destined larvae coupled with their overlapping morphospace makes it difficult to establish clear instar boundaries in *M*. *pharaonis*. Next, we tested whether developmental characters could further distinguish larval caste and sex (Figure 9 and 10). Previous studies have shown that final instar worker-destined larvae of *Monomorium emersoni and M. trageri* do not develop rudiments of wing imaginal discs, while last instar gyne/queen larvae develop wing imaginal discs that become the adult wings (Favé et al. 2015, Rajakumar et al. 2018). Moreover, it was recently shown that reproductive-destined larvae develop a germline, as marked by expression of *vasa* mRNA, while workers do not (Qui et al. 2022). Here, we show that morphologically indistinguishable 1^st^ instar can be annotated as reproductive or worker-destined based on the presence or absence of wing imaginal discs as marked by *headcase* expression and/or presence or absence of larval germ cells as marked by *vasa* expression (Figure 9a-b’). 1^st^ instar larvae that possess wing imaginal discs and germ cells are reproductives, while larvae that lack wing imaginal discs and germ cells are workers. Similarly, within the final larval instar, probing larvae for *vasa* and *headcase* mRNA provides a robust tool to distinguish caste and sex (Figure 9c-e’). Where gyne and males can be distinguished based on the morphology of the gonad marked by *vasa*. Furthermore, we show that similar to other ants, morphology of the genital imaginal disc can be used to differentiate between sexes, in both live and fixed samples (Penick et al. 2013) (Figure 10a-c’’). Finally, as a proof-of-principle we used the morphology of the genital disc in live larvae to annotate final instar reproductive-destined larvae as either gyne or male (Figure 10d). These findings suggest that the majority of our previously sample larvae may have been gyne-destined.

**Figure 9.**
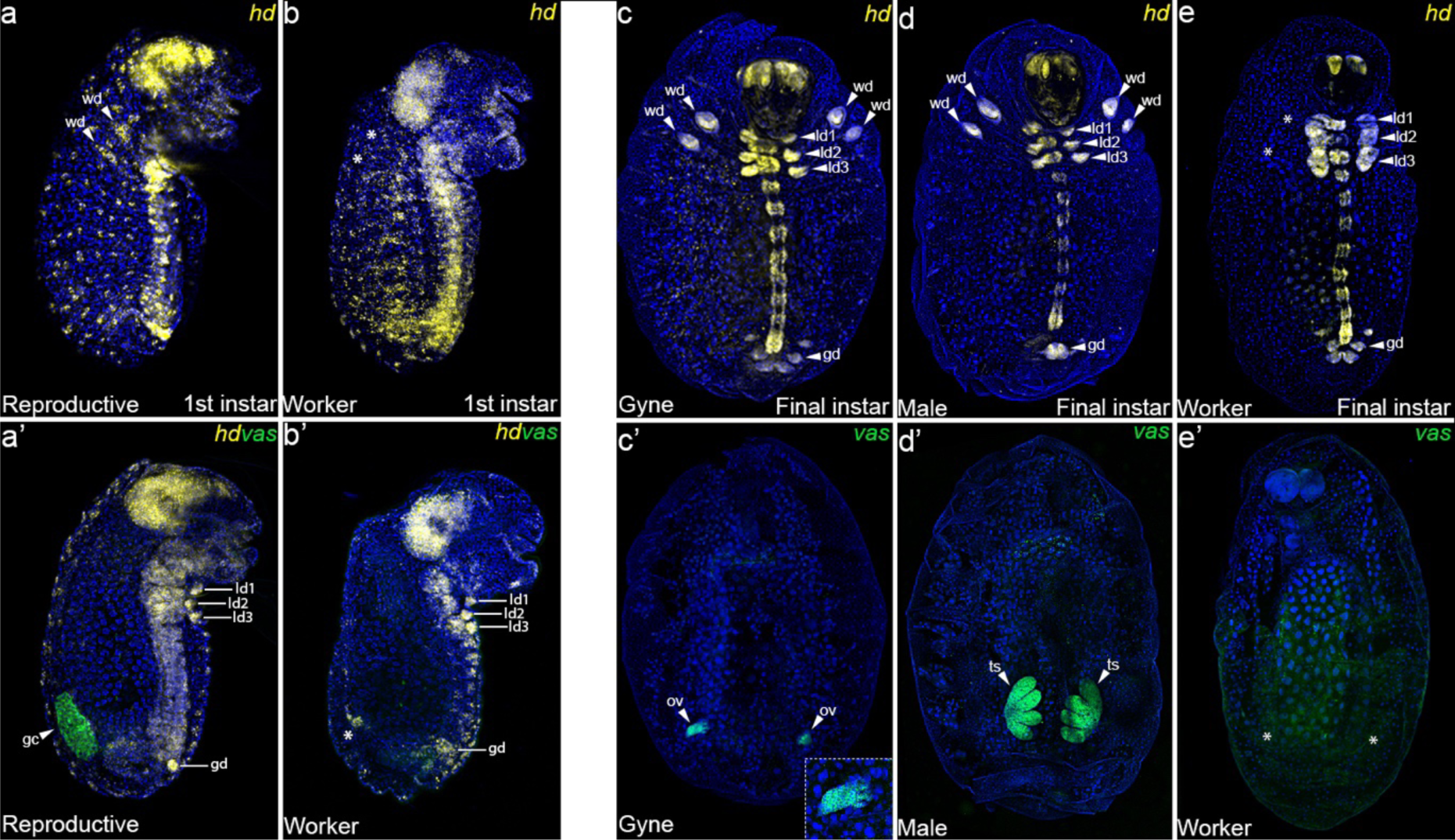
Characterization of *M*. *pharaonis* larval caste and sex using developmental markers. a-e’) *M*. *pharaonis* whole mount larvae stained with markers to distinguish caste and sex-specific developmental characters. a,b) Lateral surface of a 1^st^ instar reproductive (a) and worker (b) larvae stained with HCR probes targeting *headcase* (*hd, yellow)*), an imaginal disc marker. a) White arrows indicate the presence of wing imaginal discs (wd). b) Asterisks indicate the absence of wd. a’, b’) Sagittal plane of a 1^st^ instar reproductive (a’) and worker (b’) larvae stained with HCR probes targeting *hd*, which marks leg imaginal discs (ld1-3) and the genital imaginal disc (gd), and *vasa* (*vas*, green), which marks the larval germ cells (gc). a’) White arrow indicates the presence of germ cells. b’) Asterisk indicates absence of germ cells. c, d, e) Ventral rotation of final instar gyne (c), male (d), and worker (e) larvae stained with HCR probes targeting *hd*. c,d) White arrows indicate wd, ld1-3, and gd. e) Asterisks indicate the absence of wd. c’, d’, e’) Dorsal rotation of final instar gyne (e’), male (f’), and worker (g’) larvae stained with HCR probes targeting *vas*. c’,d’) White arrows indicate ovary (ov; c’) and tests (ts; d’). e’) Asterisks indicate the absence of ov.

**Figure 10.**
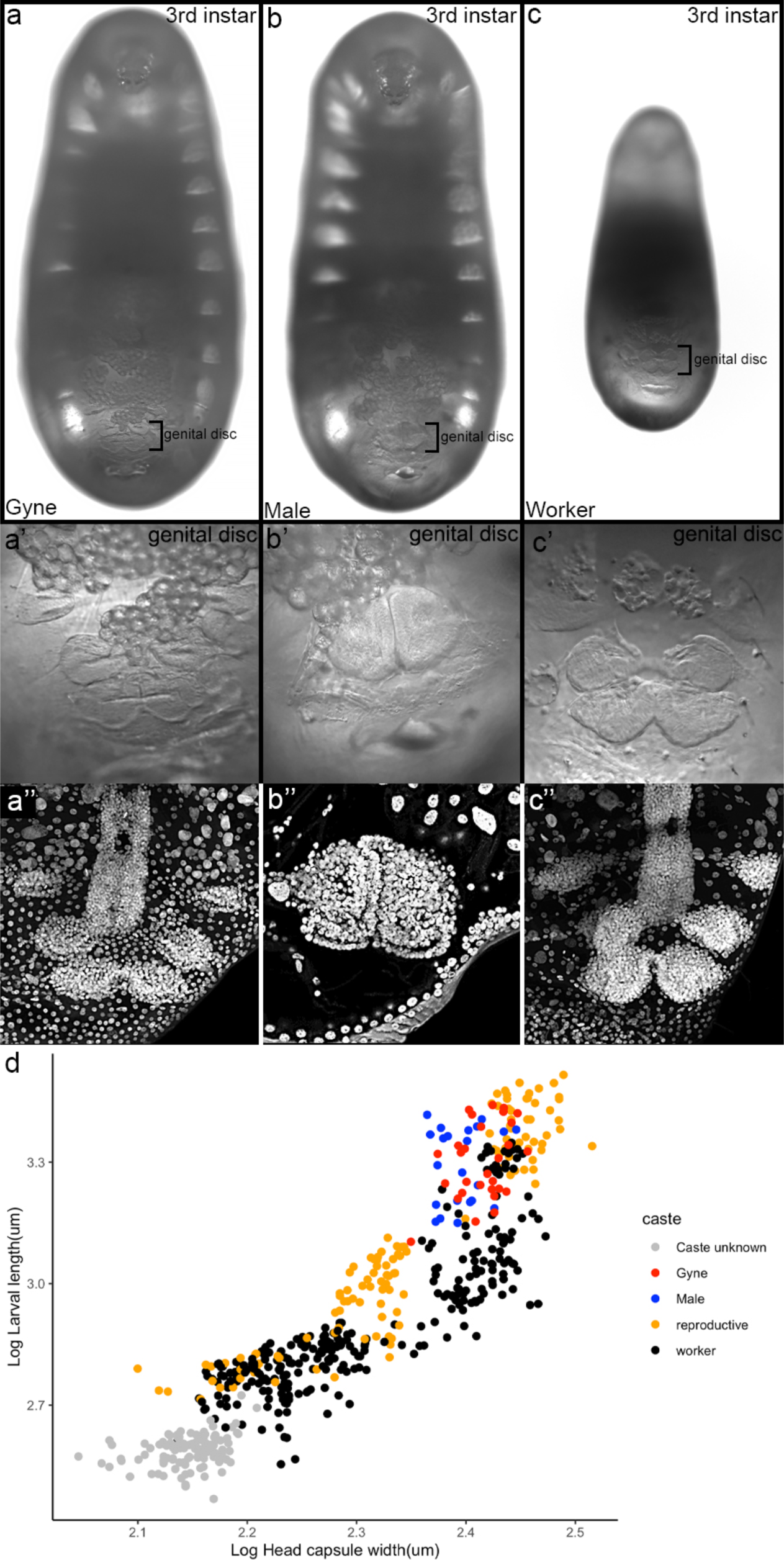
Characterization of *M*. *pharaonis* sex using genital disc morphology. a-c’) Brightfield images of gyne (a), male (b), and worker (c) final instar larvae. a’, b’, c’) High magnification images of gyne (a’), male (b’), and worker (c’) genital discs. Note the lotus shape of female castes. a’’, b’’, c’’) High magnification DAPI images of gyne (a’’), male (b’’), and worker (c’’) genital discs. All images are to scale. Plot of log Larval length by log Head capsule width of our 575 previously sampled larvae, with an additional 66 sampled last instar reproductive-destined larvae annotated based on the morphology of their genital disc using a brightfield microscope. White = Caste unknown, gold= reproductive-destined caste, black = worker-destined caste, blue = male, red = gyne.

Overall, the unique larval system that *M. pharaonis* (three types of larvae simultaneously produced) allows for the rare opportunity to chart development of multiple castes from the start of larval development through metamorphosis, knowledge of which, can provide fundamental insights toward mechanisms of caste differentiation.

### Pupal development

Pupal development for *M. pharaonis* was characterized by imaging individual worker and reproductive pupae as they aged from pupation to eclosion (Figure 11). During the pre-pupa stage, for both worker (average body length ± SD: 1.576 ± 0.070; average head width ± SD: 0.269 ± 0.006, N= 35) and reproductive (average body length ± SD: 2.630± 0.103; average head width ± SD: 0.279± 0.010, N= 24). Pre-pupal individuals can be easily distinguished from the larval stages as the cuticle undergoes extensive wrinkling. The cuticle becomes increasingly wrinkled, starting from the most posterior abdominal segments. Furthermore, pre-pupae are white, and the gut is colorless due to the meconium being expelled at the end of the final larval instar. Under our experimental conditions, pupal development took a total of 12 days, and there was no difference across castes in the time to eclosion. The length of pupal development varies greatly across ant lineages (Wheeler and Wheeler 1973, Lommelen 2003, Ishii 2005), ranging from 12-18 days in the pavement ant *Tetramorium caespitum*, 23 days in *Cryptocerus rohweri*, and up to 36 days in the ponerine ant *Pachycondyla obscuricornis*. Whether the relatively short developmental time of *M*. *pharaonis* pupae may facilitate their invasive life history remains to be determined in a broader phylogenetic framework.

**Figure 11.**
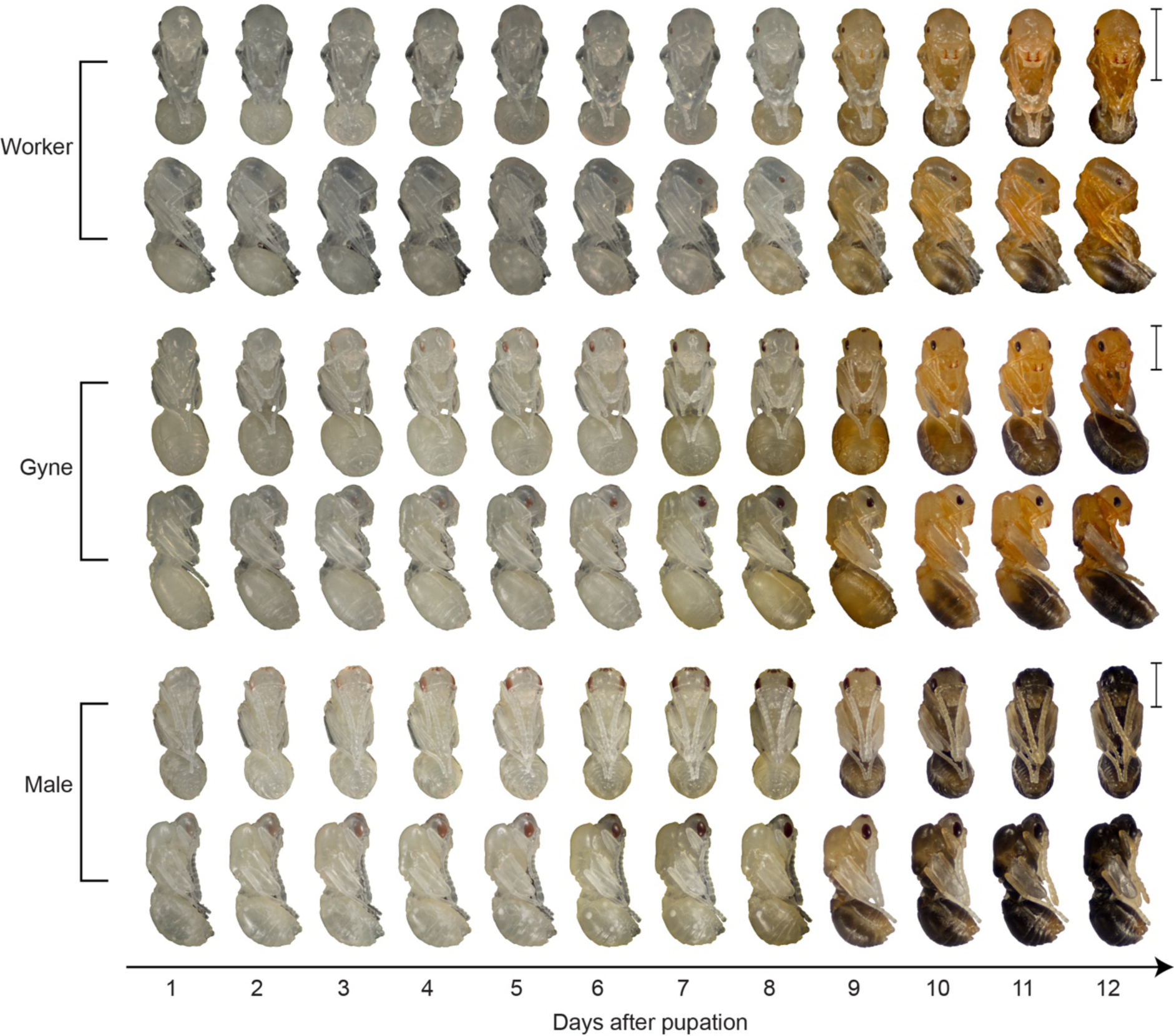
*M*. *pharaonis* pupal progression. Ventral (upper rows) and lateral (lower rows) views of individual worker (top), gyne (middle) and male (bottom) pupae for each day of pupal development until eclosion. Scale bar is 1mm.

*M. pharaonis* pupae (Figure 11) are exarate and “naked” (do not spin a silk cocoon upon pupation). Worker pupae are approximately 1.5 mm long, whereas gyne and male pupae are longer (Figure 11). Pupae of all castes become increasingly darker as they age, seemingly at the same rate. Males and females can be easily distinguished from day 1 by some morphological features. Males display bigger, ovoidal eyes, whereas gynes eyes are more rounded, and workers have significantly fewer ommatidia (Figure 12a-c). In males, antennae run almost parallel to the body for their entire length, whereas in females, antennae display a more pronounced angle, so that only the last antennal segments contact the body. Moreover, males and gynes possess three ocelli, a larger mesosoma, and two pairs of wings, which are absent in workers (Figure 12d-l). As they age, pupae acquire the characteristic coloration of the adult form: gynes have an orange-brownish head and thorax, and a black abdomen (Figure 11). Males, instead, are completely black. Pigmentation begins as early as day 2 for gyne and male pupae, and day 3 for worker pupae. The pigmentation in all castes begins in the eyes, or eyes and ocelli for gyne and male pupae. Eye pigmentation appears to occur earlier in male than gyne or worker pupae. Following the eyes, pigmentation proceeds in the more posterior segments of the abdomen, followed by the thorax for all castes.

**Figure 12.**
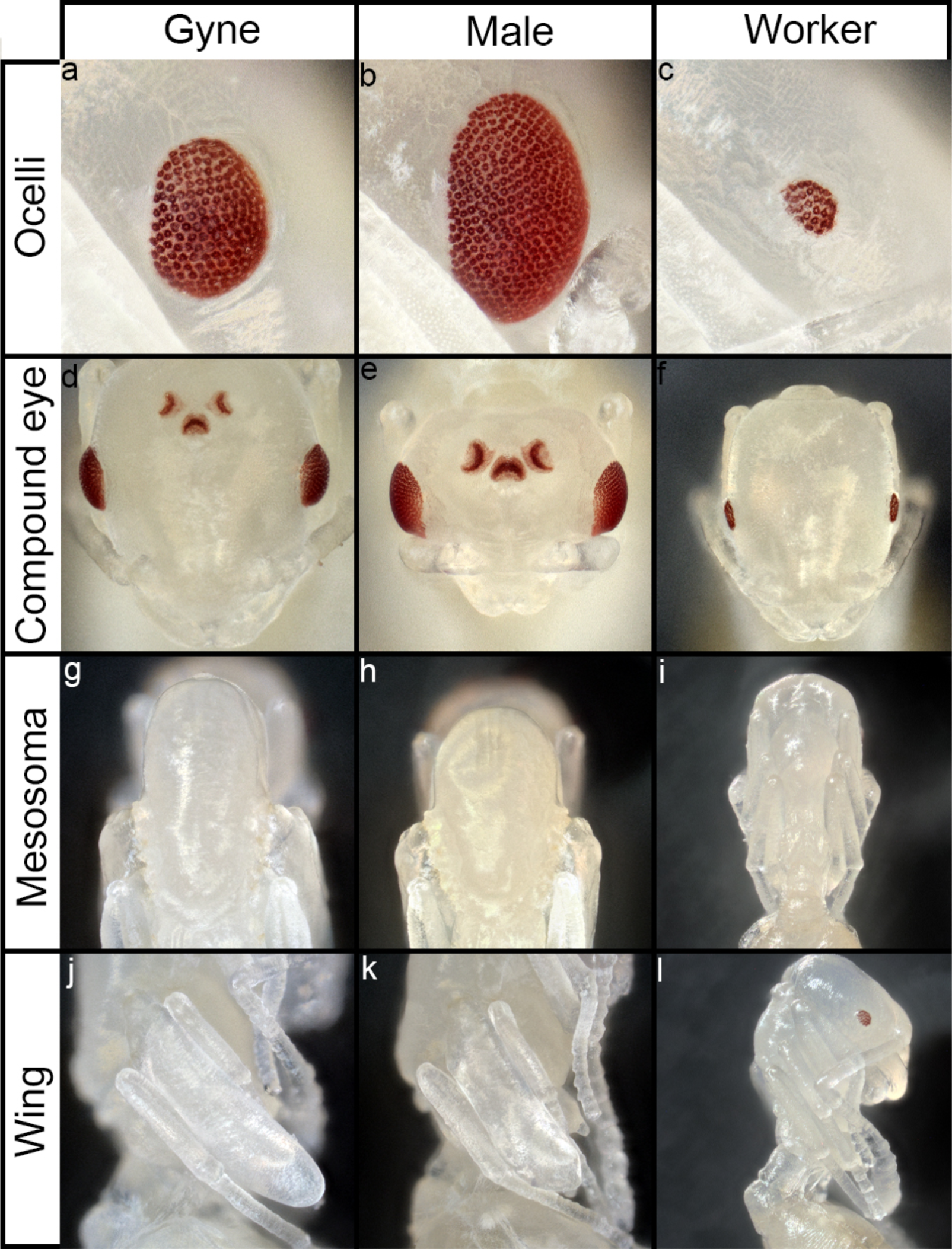
Caste and sexually dimorphic traits of *M. pharaonis*. a-l) High magnification of caste and sex-differentiated trait of gynes (left column), males (middle column), and worker (right column) pupae. a-c) Compound eyes of gynes (a), males (b), and workers (c). Note the larger and ovoidal shape of male eyes (b) and the reduction in ommatidia in worker eyes (c). d-f) Frontal view of the *M. pharaonis* head, highlight presence of absence of ocelli in gynes (d), males (e), and workers (f). g-i) Dorsal view of the *M. pharaonis* thorax, highlighting size differences in mesosoma between gynes/males (g, h) and workers (i). j-l) lateral view of the *M. pharaonis* thorax highlighting the developing wing blade in gynes (j) and males (l), while no wing blade develops in workers (l). All images across a trait are to scale.

Pupal development represents a hotspot for tissue morphogenesis (Gotoh et al. 2016). Therefore, the pupal stage will be of considerable interest for understanding the developmental underpinnings of the remarkable diversity in head, mandible, thorax, and petiole morphology. However, of all ontogenetic stages in ants, the pupal stage appears to be the least studied in terms of functional analysis of ant development and evolution. From a technical standpoint, the pupal stage provides some unique opportunities as the cuticle has not yet fully sclerotized and hardened. Simola et al. utilized the soft cuticle of the pupae to perform tissue specific injections of pharmacological inhibitors to test the molecular basis of intra-caste specific behaviors (Simola et al. 2015), while Miyazaki et al. injected pupae with RNAi targeting the yellow gene to understand sexually dimorphic body color (Miyazaki et al. 2014). Furthermore, in *Nasonia*, *Tribolium,* and *Onthophagus*, parental RNAi is routinely performed during the pupal stage prior to mating to allow for the testing of maternal effects (Lynch 2006, Lynch 2011, Miller 2012, Wasik 2012, Linz 2014). The largest technical hurdle for the adoption of such methodologies in ants is the ability to mate newly eclosed gynes and males in the lab for most ant systems. However, this is not an obstacle in *M. pharaonis* as adult males and gynes mate within their nests without the requirement of mating flights.

## Summary

We present an ant developmental staging table from egg to adult (Figure 13). Overall, development lasts approximately 45 days, with embryonic development lasting approximately 11 days, larval development approximately 22 days, and pupal development approximately 12 days. Using bright-field sand DIC microscopy, we characterized the main morphogenetic events occurring during 17 stages of embryogenesis and harmonized these stages with those of *D. melanogaster* (Figure 13, green and blue shades). Using the highly conserved germline markers *nanos*, *oskar* and Vasa, as well as live-imaging, we assessed the localization of the germ cells at different developmental stages. We uncovered two alternative types of germ-cell localization patterns in the embryo – the ‘In-phenotype and the ‘Out-phenotype,’ which represent caste differentiated embryos that give rise to sterile or fertile larvae and adults. Furthermore, using SEM, light microscopy, and morphometric data we built on the existing literature on *M. pharaonis* larval development (Alvares et al. 1993; Berndt and Eichler 1987). While we similarly identified the previously described three larval instars, our analyses further revealed a putative first instar reproductive-destined larvae. Moreover, we characterized developmental and anatomical markers to differentiate larval caste and sex, which we hope will facilitate future worker on *M*. *pharaonis* caste differentiation. Finally, we described pupal stages of *M. pharaonis* castes and put forward that morphogenetic mechanisms during pupal development have been largely understudied (Figure 13, orange shade). We end with the hope that this table will not only facilitate the use of *M*. *pharaonis* as a model for the *eco*-*evo*-*devo* of social insects but will serve as a blueprint for the generation of future developmental tables of other ant species to capture the remarkable diversity across the ants.

**Figure 13.**
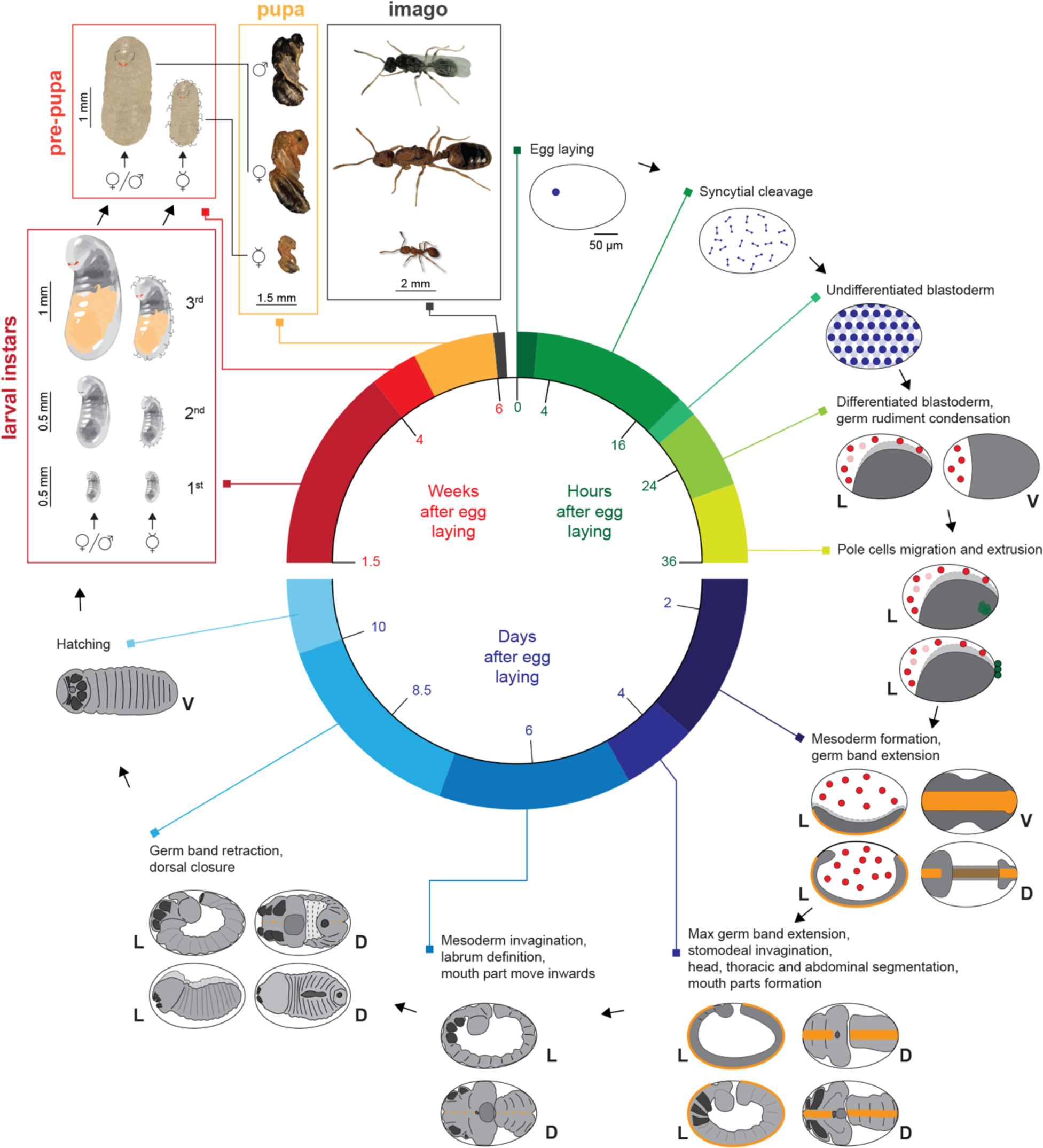
Summary diagram of *M*. *pharaonis* life cycle. Green shades indicate hours after egg laying. Blue shades indicate days after egg laying. Red to Orange indicate weeks after egg laying. Grey indicates imago. L = lateral, D = dorsal, V = ventral.

## Acknowledgements

We would like to thank the Abouheif lab for comments on the manuscript and R. Rajakumar for his advice on larval instar analyses. We would like to thank AL Price for her advice for troubleshooting larval HCR staining protocols. We also thank McGill University’s Integrated Quantitative Biology Initiative (IQBI) and Advanced BioImaging Facility (ABIF) for imaging support. This work was supported by a HFSP Long-Term Fellowship (LT0053/2022-L) to AR, a Lundbeck Foundation Grant (R190-2014-2827) to G.Z, a NSERC Discovery Grant (Canada) to E.A.

## Author contributions

LP, AR, EA and GZ conceived the project. AR, LP, RSL, GZ, and EA designed experiments. AR, LP, RL, RSL and JKLF performed experiments. AR, LP, and AMR matched ant embryonic descriptions to *Drosophila*. LP, RL, RSL, JKLF, AR and AVC collected and/or analyzed larval instar data. AR, LP, GZ, and EA wrote the manuscript. AR and LP contributed equally.

## Supplemental files

**Supplemental table 1.** Raw measurements that make up the instar plots of Figure 8 and the reproductive final instar in Figure 9.

**Supplemental video 1. Live imaging of *M*. *pharaonis embryogenesis.*** AVI, Live imagining spans from egg deposition (st1) through gastrulation and germ band extension (stage 9).

**Supplemental video 2. Live imaging of *M*. *pharaonis* ‘In’-phenotype embryos.** MP4, Live imagining of an ‘In’-phenotype embryo that spans from cellular blastoderm formation (stage 5) through the start of gastrulation (stage 6).

**Supplemental video 3. Live imaging of *M*. *pharaonis* ‘Out’-phenotype embryos.**MP4, Live imagining of an ‘Out’-phenotype embryo that spans from cellular blastoderm formation (stage 5) through the start of gastrulation (stage 6).

